# Cascading Epigenomic Analysis for Identifying Disease Genes from the Regulatory Landscape of GWAS Variants

**DOI:** 10.1101/859512

**Authors:** Bernard Ng, William Casazza, Nam Hee Kim, Chendi Wang, Farnush Farhadi, Shinya Tasaki, David A. Bennett, Philip L. De Jager, Christopher Gaiteri, Sara Mostafavi

**Affiliations:** Department of Statistics and Department of Medical Genetics, University of British Columbia, Vancouver, British Columbia, Canada, V6T 1Z4; Centre for Molecular Medicine and Therapeutics, Vancouver, British Columbia, Canada, V5Z 4H4; Department of Computer Science, University of British Columbia, Vancouver, British Columbia, Canada, V6T 1Z4; Rush Alzheimer’s Disease Center, Rush University Medical Center, Chicago, Illinois, USA, 60612; Center for Translational & Computational Neuroimmunology, Department of Neurology and the Taub Institute for Research on Alzheimer’s Disease and the Aging Brain, Columbia University Irving Medical Center, New York, NY, USA, 10027; Paul G. Allen School for Computer Science and Engineering, University of Washington, Seattle, WA 98195

## Abstract

The majority of genetic variants detected in genome wide association studies (GWAS) exert their effects on phenotypes through gene regulation. Motivated by this observation, we propose a multi-omic integration method that models the cascading effects of genetic variants from epigenome to transcriptome and eventually to the phenome in identifying target genes influenced by risk alleles. This cascading epigenomic analysis for GWAS, which we refer to as CEWAS, comprises two types of models: one for linking *cis* genetic effects to epigenomic variation and another for linking *cis* epigenomic variation to gene expression. Applying these models in cascade to GWAS summary statistics generates gene level statistics that reflect genetically-driven epigenomic effects. We show on sixteen brain-related GWAS that CEWAS provides higher gene detection rate than related methods, and finds disease relevant genes and gene sets that point toward less explored biological processes. CEWAS thus presents a novel means for exploring the regulatory landscape of GWAS variants in uncovering disease mechanisms.

**Summary:** The majority of genetic variants detected in genome wide association studies (GWAS) exert their effects on phenotypes through gene regulation. Motivated by this observation, we propose a multi-omic integration method that models the cascading effects of genetic variants from epigenome to transcriptome and eventually to the phenome in identifying target genes influenced by risk alleles. This cascading epigenomic analysis for GWAS, which we refer to as CEWAS, combines the effect of genetic variants on DNA methylation as well as gene expression. We show on sixteen brain-related GWAS that CEWAS provides higher gene detection rate than related methods, and finds disease relevant genes and gene sets that point toward less explored biological processes.

## Introduction

Genome wide association studies (GWAS) have discovered tens of thousands of common genetic variants (SNPs) associated with complex traits and disease susceptibility^1^. Identifying the targeted genes of these SNPs is critical for translating raw GWAS findings into disease mechanisms. Yet, the majority of GWAS SNPs lie in non-coding regions^2^, thus determining the genes through which these SNPs act remains challenging. The conventional approach is to apply univariate association analysis to map each SNP to its target gene based on the correlation between SNP dosages and gene expression levels (i.e. expression quantitative trait loci (eQTL) studies)^3^. This approach is often followed up by applying statistical techniques that test the probability of colocalization between GWAS SNPs and molecular QTLs in finding causal SNPs^4–7^. More recent approaches for finding disease-associated genes are converging toward using sparse models to select the combination of *cis* SNPs near each gene that together are predictive of gene expression levels^8,9^. A few studies have begun to investigate epigenomic modifications by combining the effects of GWAS SNPs on phenotypes and DNA methylation (mQTLs), in addition to gene expression (eQTLs)^10–12^. Along these lines, the use of epigenomic annotations to guide selection of expression-predictive SNPs has also been proposed^13^. Further, a recent approach attempts to go beyond modeling *cis* effects by additionally incorporating *trans* SNPs associated with epigenomic mediators of gene expression^14^.

While the importance of combining multiple omics data types is increasingly recognized for gene prioritization, most existing methods do not capture the cascading mechanism through which regulatory SNPs eventually act on phenotypes. To better trace the functional consequence of genetic variants, we propose a multi-omic integration method that mirrors the biophysics of SNP effects on nearby epigenomic elements^15^, which in turn impact gene expression and ultimately phenotypes. Our method captures this cascading mechanism by coupling two types of prediction models: one that links *cis* genetic effects to epigenomic variation, and another that links *cis* epigenomic variation to gene expression, which is analogous to using two-stage regression to model mediation effects of the epigenome on gene expression with SNPs being the instrumental variables. Here, we focus on *cis* effects, since *cis* effects tend to be more replicable than their *trans* counterparts^16^. Applying these models in cascade to GWAS summary statistics generates gene level statistics that reflect genetically-driven epigenomic effects on a given phenotype. We thus refer to our method as <u>c</u>ascading <u>e</u>pigenomic analysis for G<u>WAS</u> (CEWAS).

To test CEWAS, we first build the respective prediction models using imputed genotype^17^, DNA methylation (DNAm)^18^, and RNAseq^19^ data from the Religious Orders Study and Rush Memory and Aging Project (ROSMAP)^20,21^, which is the largest brain tissue dataset with all three data-types. We then apply CEWAS to sixteen well-powered, brain-related GWAS^22–36^, and compare it against the closest state-of-the-art methods, namely MetaXcan^9^ and EpiXcan^13^. We show that CEWAS achieves higher gene detection rate, and is able to identify disease relevant genes and gene sets that are missed by the contrasted methods. CEWAS thus provides a novel multi-omic means for inferring disease mechanisms from the regulatory landscape of GWAS variants.

## Results

### Data and model building

CEWAS builds upon MetaXcan^5^ by additionally modeling the cascading effects of SNPs on gene expression through epigenomic marks. In particular, CEWAS entails learning a set of models for predicting DNAm levels from genotype data and a set of models for predicting gene expression from *predicted* DNAm levels (Fig 1a, Methods). The former set of models are applied to GWAS summary statistics to generate epigenomic level statistics, which are then combined using the latter set of models on a gene-by-gene basis. CEWAS is thus analogous to two-stage regression used in instrumental variable analysis in modeling the mediation effects of CpGs on gene expression with SNPs being the instrumental variables. To build the prediction models, we used imputed genotype^17^, DNAm^18^, and RNAseq^37^ data from ROSMAP^20,21^, which were all derived from dorsolateral prefrontal cortex (DLPFC) tissue of ∼700 individuals. Both gene expression and DNAm data were corrected for hidden covariates and measured technical confounders (see Methods). For each CpG, we built a DNAm prediction model using elastic net^38^ with dosage level of SNPs within ±50Kb of that CpG as covariates. Similarly, for each gene, we built an elastic net expression prediction model with *predicted* DNAm level of CpGs within ±500Kb of that given gene. Using these learned model weights, we applied CEWAS to a range of brain-related GWAS^22–36^ to find their implicated genes, as we describe next.

**Figure 1.**
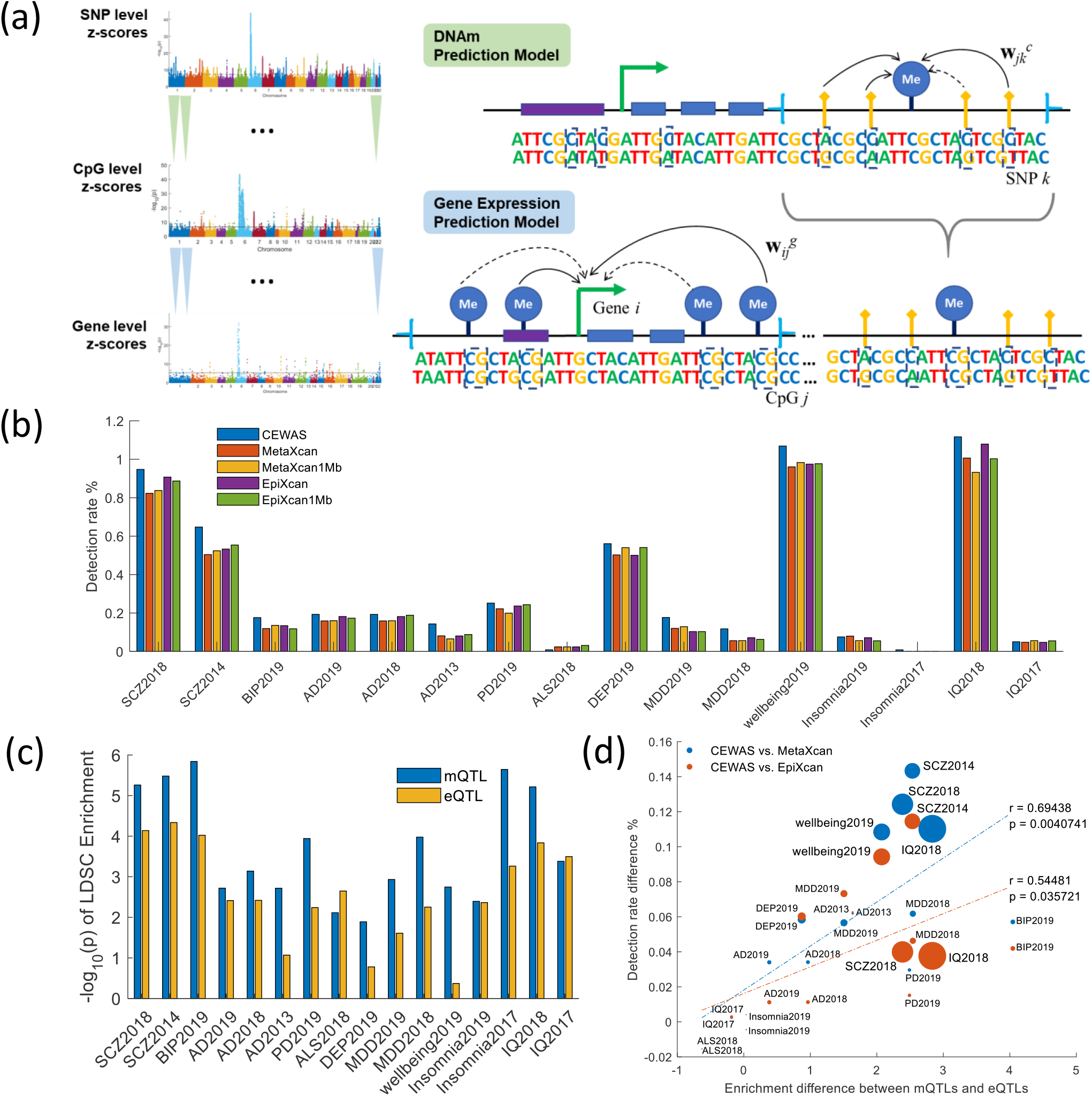
CEWAS and gene detection. (a) For each CpG *j*, a DNAm prediction model is built using dosage levels of proximal SNPs, and for each gene *i*, a gene expression prediction model is built using predicted DNAm levels of proximal CpGs. Elastic net is used for model learning, where solid and dotted lines pictorially indicate non-zero and zero weights for the corresponding variable pairs. These models are applied in cascade to GWAS summary statistics to estimate gene level z-scores. (b) Bars show the detection rate (defined as the percentage of tested genes declared significant) for the examined GWAS. (c) Bars show the log p-value of enrichment of mQTLs and eQTLs assessed by LDSC for each GWAS. The baseline SNP sets in LDSC corresponding to various regulatory attributes are included as background. (d) Difference in mQTL vs. eQTL log p enrichment (from Figure 1c) plotted against difference in detection rate between CEWAS and a contrasted method (color of the points indicate which method is being contrasted with CEWAS). The size of a point is proportional to the polygenecity of the corresponding GWAS (approximated by the ratio of the number of reported GWAS loci over the sample size). As shown, enrichment for mQTLs over eQTLs significantly correlates with improvement in detection achieved by CEWAS over contrasted methods (p<0.05).

### Prioritizing disease-associated genes

We applied CEWAS to sixteen well-powered, brain-related GWAS^22–36^ (Table S1), and compared it against MetaXcan and EpiXcan (which CEWAS is built upon) in terms of gene detection rate, defined as the proportion of expressed genes identified as significant at an α of 0.05 with Bonferroni correction. MetaXcan and EpiXcan models were built using the same input SNPs and expression dataset as CEWAS. As shown in Fig 1b (the number of tested and detected genes summarized in Table S2), CEWAS achieved higher detection rate than MetaXcan (p=0.00074, Wilcoxon sign rank test across GWAS), which shows the benefits of integrating DNAm information. This trend remained for MetaXcan models built using SNPs within the typical window of ±500Kb from transcription starting site (TSS) of genes (denoted as MetaXcan1Mb). CEWAS also achieved higher detection rate than EpiXcan (p=0.00088), which demonstrates that using DNAm data to incorporate additional models, instead of weighting SNPs by epigenomic annotation, increases detection rate. The same trend was observed with EpiXcan models built using SNPs within ±500Kb from TSS of genes (denoted as EpiXcan1Mb). Since SNPs might be shared between models of spatially proximal genes, we further estimated detection rate based on the number of *distinct* signals (see Methods), and confirmed that the higher detection rate attained by CEWAS remains to hold with this analysis (Fig S1).

A natural question is what gave rise to CEWAS’s higher detection rate. Since CEWAS does not directly model the associations between SNP dosages and gene expression levels, one would expect its accuracy in predicting gene expression (R^2^) to be lower than MetaXcan and EpiXcan, which was indeed what we observed (Fig S2-S3). This lower R^2^ rules out higher predictive accuracy as the reason for CEWAS’s higher detection rate. Instead, we hypothesized that the higher detection rate arises from the type of SNPs selected by CEWAS. Specifically, SNPs selected by CEWAS are largely mQTLs by construction, whereas SNPs selected by MetaXcan and EpiXcan are primarily eQTLs. To test this hypothesis, we estimated the partitioned heritability of mQTLs against eQTLs using linkage disequilibrium score regression (LDSC)^39^. We examined mQTLs and eQTLs derived from the ROSMAP data^15^, but restricted to those where the mQTL SNPs are within a 100Kb window from CpGs (i.e. matching the window size used for CEWAS), and the eQTL SNPs are within a 1Mb window from TSS of genes (i.e. typical window size for eQTL analysis). The baseline SNP sets in LDSC corresponding to various regulatory attributes were included as background. As shown in Fig 1c, the enrichment for mQTLs is higher than for eQTLs (p<0.0084, Wilcoxon sign rank test across GWAS), and the same trend holds when we matched the window size of eQTLs to that of mQTLs, i.e. 100Kb, as well as matching the number of mQTL SNPs to the number of eQTL SNPs (Fig S4). Indeed, in support of our hypothesis, this higher enrichment for mQTLs over eQTLs was found to significantly correlate with the increase in detection rate achieved by CEWAS over MetaXcan and EpiXcan (Fig 1d). Also, mQTLs tend to be more replicable than eQTLs (Fig S5), hence CEWAS would presumably be more robust than MetaXcan and EpiXcan in selecting relevant SNPs across GWAS, which could further explain CEWAS’s higher detection rate. In fact, similar trends of higher GWAS enrichment and reproducibility with mQTLs over eQTLs have also been seen in blood^40^, which supports our observations, though whether these trends are generalizable to other tissues requires further investigation.

The next question is whether modeling DNAm mediated effects on GWAS phenotypes alone, i.e. without using expression data, is already adequate to attain higher detection rate. To answer this question, we assessed DNAm mediated effects on GWAS phenotypes in three ways. First, we directly examined results from models used in the first stage of CEWAS, i.e. MetaXcan models built with each CpG taken as the response and SNPs within ±50Kb from that CpG as predictors. Detection rate of these DNAm MetaXcan models, defined as the number of detected CpGs among tested CpGs, is significantly lower than CEWAS (p=0.0004), MetaXcan (p=0.0013), and EpiXcan (p=0.0006) based on Wilcoxon sign rank test across GWAS (Fig S6, Table S2). Second, we mapped CpGs to genes without using expression data, by taking the DNAm MetaXcan p-value of the closest CpG of each gene as the p-value of that gene. The gene level detection rate is significantly lower than CEWAS (p=0.0004), MetaXcan (p=0.0003), and EpiXcan (p=0.0004). Third, we mapped CpGs to genes by taking the DNAm MetaXcan p-value of the CpG with the largest R^2^ in terms of DNAm prediction among CpGs within ±500Kb from each gene as the p-value of that gene. The detection rate is lower than CEWAS (p=0.1331) and EpiXcan (p=0.3808), and higher than MetaXcan (p=0.4235) but not significant. Based on these comparisons, mapping CpGs to genes without using expression data does not provide the same detection gain as CEWAS. Also, the highlighted genes are different from the expression-based models (i.e. CEWAS, MetaXcan, and EpiXcan) as assessed using cross-method area under the receiver operating characteristic curve (AUC). Specifically, for each GWAS, we ranked genes based on p-values from one of the expression-based models, took the top 1% of genes as reference, and estimated AUC with gene level p-values of the DNAm-based models (see replication AUC under Methods for details on AUC estimation). The cross-method AUC between DNAm models with mapping based on the closest CpG to each gene and expression-based models are 0.75±0.05, 0.70±0.04, and 0.70±0.05 for CEWAS, MetaXcan, and EpiXcan, respectively. As for mapping based on the max R^2^ CpG of each gene, the cross-method AUC are 0.65±0.05, 0.64±0.04, and 0.65±0.04 for CEWAS, MetaXcan, and EpiXcan. Hence, while mapping based on max R^2^ CpG resulted in higher detection rate than mapping based on closest CpG, the cross-method AUC against expression-based models are lower, and these AUC are much lower than cross-method AUC between expression-based models, which are in the range of 0.93 to 0.99. In fact, the two non-expression-based CpG-to-gene mapping approaches themselves highlight quite different genes, with cross-method AUC being only 0.63±0.05.

### Calibration test

Since CEWAS is built upon MetaXcan, which entails an approximation in its z-score estimation^5^, the increase in detection rate could potentially be due to artificial z-score inflation inherent in CEWAS’s mathematical formulation. To verify that the mathematical formulation is not the reason behind CEWAS’s higher detection rate, we performed simulations to test whether z-scores estimated by CEWAS are calibrated (see Methods). As shown in Fig 2, with null input z-scores having LD structure matched to the SNP covariance of the CEWAS models, CEWAS correctly returned null z-scores as output, which verifies that CEWAS z-scores are indeed calibrated when the input data match the model assumptions. However, in practice, the LD structure of GWAS genotype data might not match CEWAS’s model covariance. To test the effect of LD mismatch, we used the LD structure estimated from the 1000 Genome phase 3 genotype data (European population) to generate null input z-scores. The resulting CEWAS z-scores have standard deviation close to or less than 1 for majority of the genes except for some outliers, but the number of outlier genes is similar to MetaXcan (Fig 2). Thus, CEWAS’s higher detection rate is unlikely due to more false positive detections compared to MetaXcan and EpiXcan. Nonetheless, this result highlights the importance of LD matching for applying CEWAS (as well as MetaXcan and EpiXcan). We also note that CEWAS requires an estimate of the covariance between CpGs. Such covariance must be estimated using predicted DNAm levels since only the genetic component of DNAm is modeled in CEWAS. If the CpG covariance is estimated from the measured DNAm data, the resulting output z-scores would be inflated.

**Figure 2.**
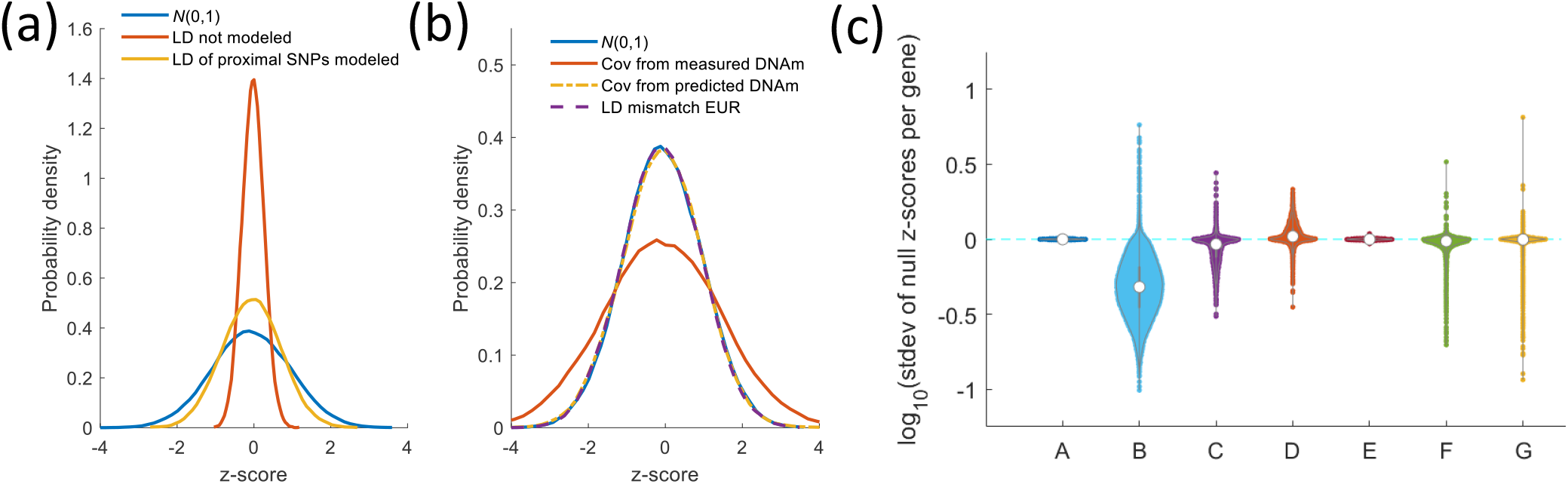
Calibration test. (a) Testing CEWAS requires modeling the LD structure of SNPs when generating null input **z**_*k*_^*s*^. Shown are probability density functions of **z**_*i*_^*g*^ for an exemplar gene derived from 10000 sets of null **z**_*k*_^*s*^ under different input settings. Red curve: **z**_*k*_^*s*^ generated from *N*(0,1). Yellow curve: **z**_*k*_^*s*^ generated with LD of only SNPs proximal to each CpG modeled. For both input settings, **z**_*i*_^*g*^ are underestimated, i.e. not matching the blue curve corresponding to *N*(0,1). (b) By modeling LD of all SNPs proximal to CpGs of a given gene when generating null **z**_*k*_^*s*^ and using CpG covariance based on predicted DNAm levels when estimating **z**_*i*_^*g*^, the resulting **z**_*i*_^*g*^ (yellow curve) are calibrated (i.e. matching blue curve), whereas using CpG covariance based on measured DNAm levels to estimate **z**_*i*_^*g*^ resulted in z-score inflation (red curve). In practice, the LD structure of GWAS genotype data might not match the SNP covariance of the CEWAS models. To test the effect of LD mismatch, we further used the LD structure estimated from the 1000 Genome phase 3 genotype data (European population) to generate **z**_*k*_^*s*^. The resulting **z**_*i*_^*g*^ are well calibrated for this exemplar gene (purple curve). (c) Standard deviation of 10000 **z**_*i*_^*g*^’s for each gene *i* shown across all genes. A: **z**_*i*_^*g*^ drawn from *N*(0,1). B: input **z**_*k*_^*s*^ drawn from *N*(0,1). C: **z**_*k*_^*s*^ with LD of SNPs proximal to each CpG modeled. D and E: **z**_*k*_^*s*^ with LD of all SNPs modeled and using CpG covariance based on measured and predicted DNAm levels, respectively, when estimating **z**_*i*_^*g*^. Standard deviation of **z**_*i*_^*g*^ in E is ∼1 for all genes, confirming that CEWAS produces calibrated **z**_*i*_^*g*^ when LD of **z**_*k*_^*s*^ is matched to CEWAS’s model covariance. F and G: **z**_*k*_^*s*^ with LD estimated from 1000 Genome data, and **z**_*i*_^*g*^ estimated by CEWAS and MetaXcan, respectively. Standard deviation of CEWAS **z**_*i*_^*g*^ are close to or less than 1 for majority of genes except for some outliers, and the number of outlier genes is similar to MetaXcan.

### Replication of CEWAS models

To assess the generalizability of CEWAS, we built another set of prediction models using genotype and RNAseq data generated from DLPFC tissues of 592 subjects in the CommonMind Consortium (CMC) study^41^ in combination with mQTLs derived from DLPFC DNAm data of 526 subjects in the DevMeth study^42^ (see Methods), and applied these models to the same set of GWAS^22–36^ as with the ROSMAP models. We needed to combine datasets from different studies since, to the best of our knowledge, no current brain tissue-based studies other than ROSMAP collected all three data-types from the same individuals. As the replication metric, we used AUC, as computed by ranking the genes of each GWAS based on p-values estimated with the ROSMAP models, taking the top 1% of genes as the reference, and estimating AUC with p-values from the CMC models (see Methods). We opted to use the top 1% of genes since less than a handful of genes are statistically significant for some of the GWAS, which might be too few for robust AUC estimation. CEWAS achieved an average AUC of 0.76±0.04 (Table S3), which is significantly higher than the chance level AUC of 0.5 as confirmed with permutation test. For comparison, we repeated this analysis for MetaXcan and EpiXcan with their corresponding models built using ROSMAP vs. CMC data, which attained average AUC of 0.80±0.05 and 0.87±0.03, respectively. Using the top 1% of genes detected by the CMC-based models as reference resulted in the same trend, with average AUC of 0.80±0.05, 0.82±0.04, and 0.86±0.03 attained by CEWAS, MetaXcan, and EpiXcan, respectively. As one would expect, the replication AUC of CEWAS is lower than MetaXcan and EpiXcan since replication of CEWAS requires combining two data sources (CMC and DevMeth) and matching more variables across datasets. Indeed, following up on this result, we assessed cross-method AUC (Table S3) by using the top 1% of genes from CMC-based MetaXcan and EpiXcan models as reference and p-values from ROSMAP-based CEWAS models for AUC estimation. The cross-method AUC are 0.83±0.05 and 0.87±0.04 with MetaXcan and EpiXcan as reference, respectively, which are on par with replication AUC of MetaXcan and EpiXcan. Thus, the need to combine two disparate data sources for replicating CEWAS seems to be the key reason to its lower replication AUC.

### Genomic correlation between GWAS

As another form of replication, we examined the correlation between CEWAS z-scores across the sixteen GWAS^22–36^ (Fig 3a), an idea that has been used for studying relationships between complex traits^43^. For the *same* phenotype examined in separate GWAS samples, the genomic correlations of CEWAS z-scores range from 0.44 to 0.93 (Fig 3b). For well-powered GWAS pairs of the same phenotype, coherent z-scores were observed (r>0.8, Figure 3b), while the cases of lower genomic correlation could partly be explained by large differences in sample size between the GWAS pairs (Fig 3c). When looking at genomic correlation between different phenotypes, for schizophrenia and bipolar disorder, a correlation of 0.4 between CEWAS z-scores was found, which recapitulates previous findings^44^. Both disorders show correlations of ∼0.17 with depression. CEWAS found four genes that are common across these disorders (Fig 3d): *GNL3, SPCS1, TMEM110*, and *MCHR1*. Fittingly, overexpression of *GNL3* was previously shown to reduce the density of mushroom dendritic spines in rats, which might relate to dendritic spine pathology observed across patients with schizophrenia, bipolar disorder, and depression^45^. Similar to *GNL3, SPCS1* and *TMEM110* also lie in the 3p21.1 region, and their expression levels were previously shown to be associated with risk variants of schizophrenia, bipolar disorder, and depression^45^. In fact, CEWAS also detected *GNL3, SPCS1*, and *TMEM110* for intelligence with z-scores having opposite signs compared to schizophrenia (Fig S7), which aligns with how risk alleles in these loci were previously shown to correlate with lower cognitive test scores^45^. Results obtained with MetaXcan and EpiXcan display the same trend, but only CEWAS found *MCHR1* to be common across schizophrenia, bipolar disorder, and depression. Specifically, while *MCHR1* was also found to be significant for schizophrenia using MetaXcan and EpiXcan, their p-values for bipolar disorder and depression are an order of magnitude higher than the Bonferroni threshold. Fittingly, *MCHR1* and limbic regions, such as amygdala and hippocampus in which *MCHR1* is expressed, are modulators of stress response^46^. Also, although the genomic correlation between Alzheimer’s disease (AD) and Parkinson’s disease (PD) is only 0.05 (which matches the low genetic correlation of 0.08 observed in a previous study^47^), some shared genes were found (Fig 3d). In particular, all contrasted methods found *C16orf93* and *KAT8* to be common between AD and PD, but only CEWAS additionally found *PRSS36* and *ZNF668*. These genes were also found by MetaXcan and EpiXcan for AD but not PD, with p-values being an order of magnitude higher than the Bonferroni threshold. Knockdown of *ZNF668* has been shown to impair homologous recombination DNA repair^48^. Both *PRSS36* and *ZNF668* belong to the *KRAB-ZFP* cluster, which is involved with cell proliferation^49^, hence aligns with impaired hippocampal neurogenesis being a potential mechanism underlying memory deficits in AD^50^ and impaired precursor cell proliferation due to dopamine depletion in PD^51^.

**Figure 3.**
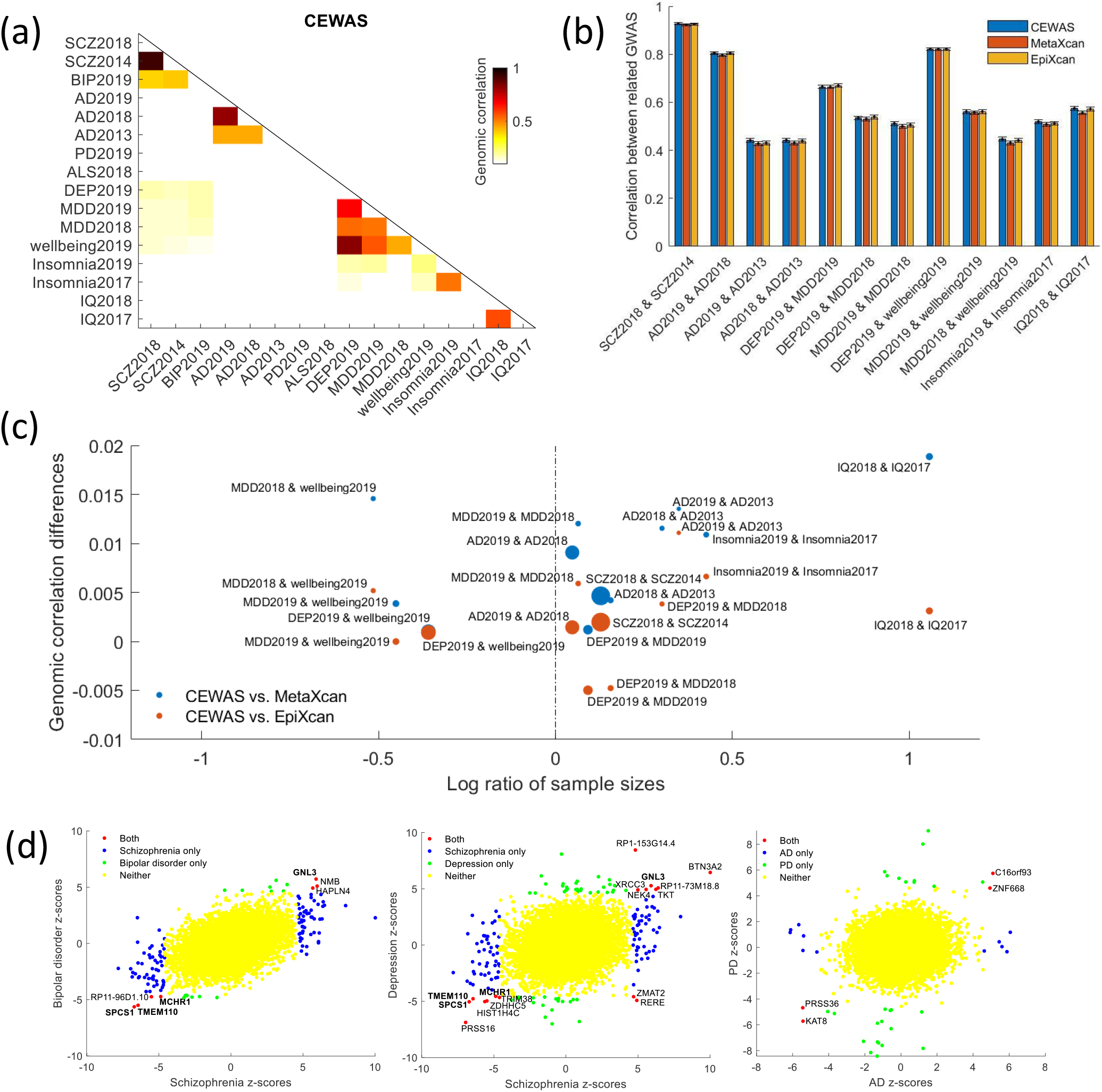
Genomic correlation across GWAS. (a) Correlation between CEWAS-derived z-scores for all GWAS pairs shown (values clipped at 0.1 to highlight higher correlation). (b) Correlations between gene level z-scores for GWAS pairs of the same phenotype displayed. Note that wellbeing spectrum is partly based on depressive symptoms, hence wellbeing2019 is expected to be highly correlated with DEP2019, MDD2019, and MDD2018. (c) Dissecting results in panel b, log ratio between sample size of GWAS pairs is plotted against differences in genomic correlation between CEWAS vs. a contrasted method. The size of a dot corresponds to the correlation between CEWAS z-scores of a GWAS pair. Two trends were observed: i. higher correlations between GWAS pairs with similar sample size (dots closer to 0 on x-axis are larger), ii. CEWAS typically yields higher genomic correlation than the contrasted methods (most dots above 0 on y-axis). (d) CEWAS z-scores of schizophrenia vs. bipolar disorder, schizophrenia vs. depression, and AD vs. PD shown as scatterplots. Genes detected across schizophrenia, bipolar disorder, and depression highlighted in bold. Although AD and PD show little genomic correlation, a few shared genes were found.

### Method similarity and differential genes

We next assessed the similarity between methods by examining the correlation between their z-scores for each GWAS. We observed an average correlation of 0.74 between CEWAS and MetaXcan (Fig 4a), which mirrors the reported overlaps between eQTLs and mQTLs from the same tissue^15^. As for CEWAS vs. EpiXcan, the average correlation is also 0.74, which might seem surprising since both CEWAS and EpiXcan use epigenomic information, whereas MetaXcan does not. The reason is that the average correlation between MetaXcan and EpiXcan is 0.93, suggesting that SNPs selected by EpiXcan are largely eQTLs analogous to MetaXcan.

**Figure 4.**
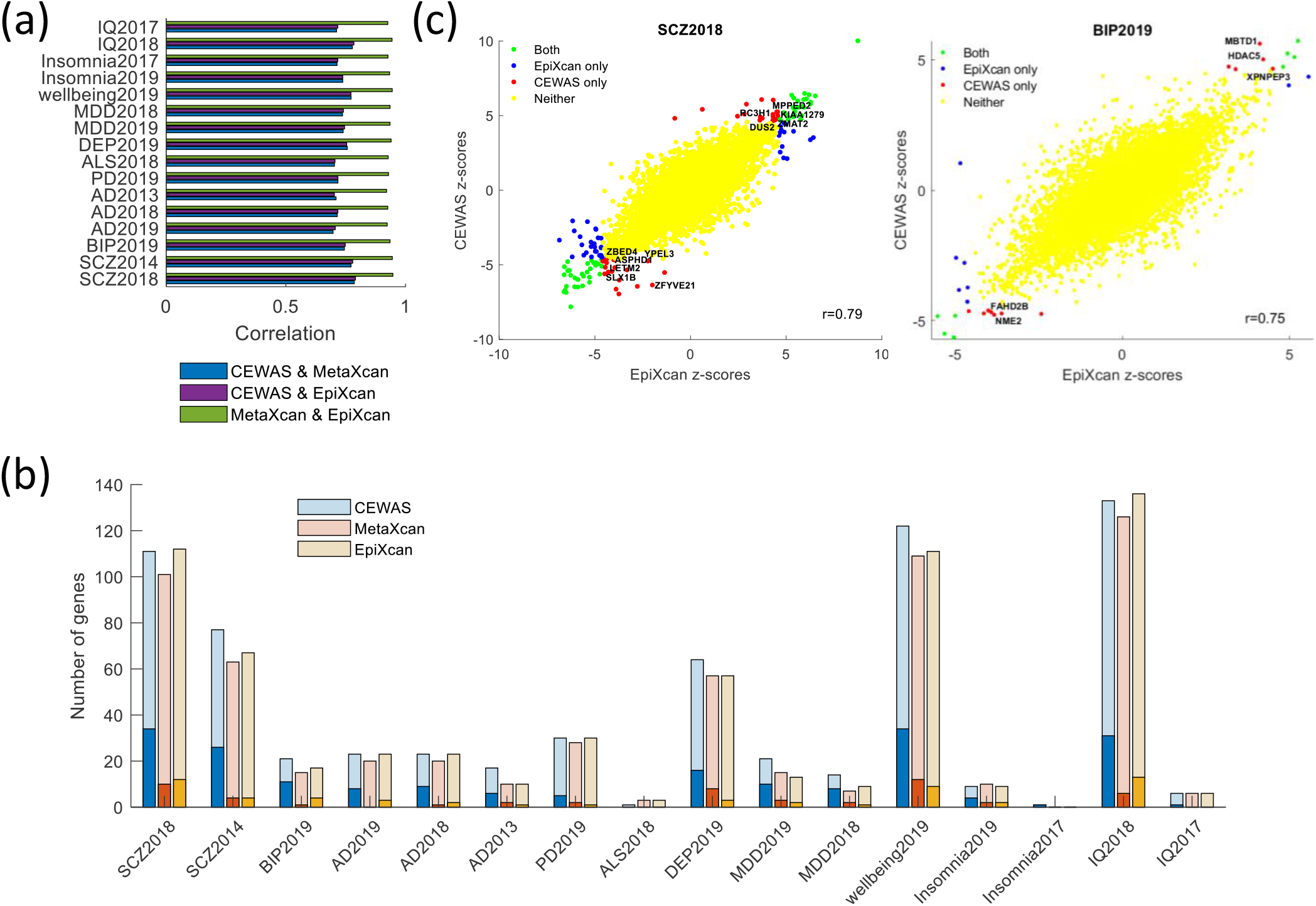
Method similarity and differential genes. (a) For each GWAS, correlation between z-scores estimated by the contrasted methods are shown. EpiXcan and MetaXcan tend to produce highly correlated z-scores. CEWAS z-scores are slightly more correlated with EpiXcan than with MetaXcan. (b) The darker shaded bars show the number of significant genes that were exclusively found by each method. The lighter shaded bars show the total number of significant genes found by each method. Only genes tested in all three methods were considered in computing the number of genes exclusively found by one method but not the other two. The number of genes exclusively found by CEWAS outnumbered MetaXcan and EpiXcan. (c) z-scores estimated by CEWAS vs. EpiXcan for SCZ2018 and BIP2019 displayed. Genes related to mitochondria dysfunction and oxidative stress exclusively detected by CEWAS but missed with MetaXcan and EpiXcan annotated.

Contrasting the significant genes detected by each method (Table S4), we observed a larger number of genes that were exclusively found by CEWAS but not by the other methods (Fig 4b). A similar trend was observed when we assessed the number of distinct signals (Fig S8). To interpret the genes only found by CEWAS, we first grouped these genes into key shared mechanisms using Mammalian phenotypes and GO terms^52^, and then searched the literature for experimental evidence of their disease relevance. For schizophrenia (Fig 4c), only CEWAS detected *DUS2, ENDOG, KIAA1279, LETM2*, and *ZMAT2*, which are related to mitochondria functions, as well as *ASPHD1, ENDOG, MPPED2, RC3H1, SLX1B, YPEL3, ZBED4, ZFYVE21*, and *ZMAT2*, which are related to metal ion-binding. Detection of these genes aligns with how the presence of reduced metal ions promotes formation of highly reactive hydroxyl radicals from mitochondrial superoxide^53^, which could increase oxidative stress, resulting in behavioral and molecular anomalies seen in schizophrenia patients^54^. Also, only CEWAS found *FAHD2B, HDAC5, MBTD1, NME2*, and *XPNPEP3* for bipolar disorder. These genes are again related to mitochondria and metal ion binding (Fig 4c), hence are linked to oxidative stress, which is a key process in the progression of bipolar disorder^55^. Among the unique genes found by CEWAS for AD, *ADAM10* plays a prominent role in the cleavage of Alzheimer’s precursor protein^56^, *ERCC1*-*XPF* endonuclease is involved with DNA repair and accelerated aging^57^, and *FAM63B* is involved with cleavage of ubiquitin and abnormal ubiquitin aggregates are often seen in AD^58^. For PD, CEWAS detected *PLEKHM1* and *TTC19. PLEKHM1* is an effector that jointly binds LC3, Rab7, and HOPS complex for lysosomal protein degradation^59^, and Rab7 has been shown to induce clearance of α-synuclein aggregates^60^. The binding of *TTC19* to mitochondrial respiratory complex III is required for *UQCRFS1* fragment clearance, deficiency of which could result in neurological impairments^61^, and oxidative stress arising from mitochondrial dysfunction has been associated with dopaminergic neuronal death in PD^62^. For depression, CEWAS uniquely detected *TNKS2* and *ZDHHC5. TNKS2* is a poly-ADP-ribosyltransferase involved in Wnt/β-catenin signaling, and β-catenin has been shown to mediate antidepressant responses^63^. *ZDHHC5* is a palmitoyl acyltransferase, and attenuated 5-HT1AR palmitoylation has been shown to induce depression-like behaviors^64^. Note that more than half of the genes exclusively found by CEWAS lie at least 100Kb away from any GWAS SNPs (Fig S9). These distal relationships are unlikely to be found using linear distance to map GWAS SNPs to target genes.

Genes only found by MetaXcan and EpiXcan also suggest interesting disease mechanisms. For instance, *DDHD2* was found for schizophrenia, which is a new candidate risk gene recently discovered based on its effects on RNA-binding protein dysregulation^65^. Also, *HSD3B7* was found for AD, which is involved with bile acid synthesis, and altered bile acid levels in relation to cognitive decline in AD has not been observed and studied until recently^66,67^. *SETD1A*, which is a schizophrenia-susceptibility gene, was interestingly also found for AD. Mutation of *SETD1A* has been shown to result in working memory deficit in mice^68^ and impaired excitatory synaptic transmission in pyramidal neurons within medial PFC^69^, which are indeed characteristics of AD.

### Gene set enrichment and brain region association

To infer the biological processes captured by CEWAS, we applied MAGMA^70^ (see Methods) to assess the enrichment for known gene sets (CNS^22^ and MSigDB^71^) and brain regions (regional expression profiles from Allen Institute^72^). In brief, the top gene sets highlighted by CEWAS point toward similar biological processes as MetaXcan and EpiXcan (Fig S10, Table S5) but with some notable differences as we describe next.

For psychiatric disorders (Fig 5a), only CEWAS highlighted CNS gene sets related to abnormal glial cell morphology and abnormal prepulse inhibition for schizophrenia (Fig 5b), bipolar disorder, and depression. Disruption of astrocytic functions arising from glial cell loss has been suggested as a potential cause for the neuronal abnormalities, such as reduced neuronal size, observed across major psychiatric disorders^73^, and decreased prepulse inhibition has been found in schizophrenia patients^74^, bipolar patients during manic state^75^, and depression patients to a lesser extent^76^. CEWAS also highlighted abnormal subventricular zone morphology for schizophrenia and depression, which aligns with how dysregulated neurogenesis has been posited as a mechanism for aberrant pattern separation and abnormal reward processing in schizophrenia and depression patients^77^. Fittingly, based on the Allen Institute data (Table S6), CEWAS highlighted the lateral nucleus of the amygdala for schizophrenia and bipolar disorder, parahippocampal gyrus, the CA4 field of the hippocampus, and basolateral nucleus of the amygdala for schizophrenia and depression, which aligns with the roles of these brain regions in emotion and reward processing as well as in startle reflex^78^ as used for prepulse inhibition. CEWAS also highlighted the dentate gyrus for schizophrenia and depression, which is the other main brain region, in addition to subventricular zone, where neurogenesis persists in adulthood^79^. Further, CEWAS highlighted raphe nuclei of medulla, reticular nucleus, and other thalamic nuclei for schizophrenia and depression. The raphe nuclei is the origin of most forebrain serotonin innervation, which aligns with the abnormal serotonin levels seen in psychiatric patients and how serotonin is used for treating psychiatric disorders^80^. The thalamic reticular nucleus (TRN) is largely composed of GABAergic neurons that express parvalbumin, and the reduction of these neurons in the TRN of psychiatric patients has been suggested to induce attentional, cognitive, and emotional deficits^81^. Moreover, CEWAS highlighted a CNS gene set related to dopaminergic neuron morphology for schizophrenia, which aligns with the dopamine hypothesis^82^, and genes found by CEWAS for depression are highly expressed in the ventral tegmental area, which is one of the main dopaminergic areas in the brain with projections to amygdala and hippocampus for emotion and reward processing as discussed, and reduced dopamine has been linked to depressed symptoms^83^.

**Figure 5.**
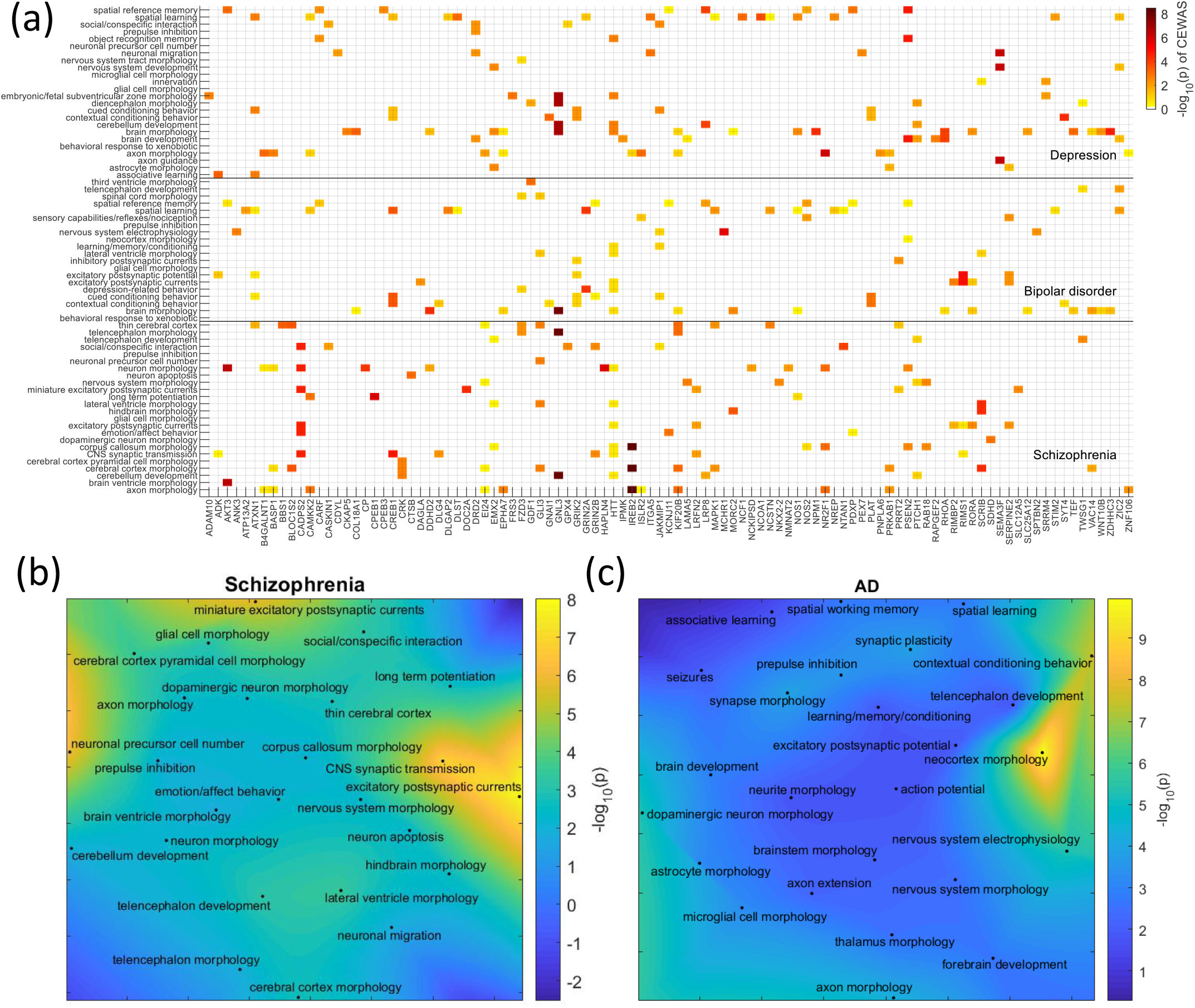
Gene set enrichment. (a) Top CNS gene sets enriched for schizophrenia, bipolar disorder, and depression as found by CEWAS displayed against genes belonging to at least one of these enriched gene sets and have CEWAS p<0.05 in at least one of the three psychiatric disorders. The color reflects the level of association between a given gene and a disorder. For each gene set, only genes belonging to that gene set have their –log_10_(p) displayed, with the rest of the genes masked to zero. (b) Top 25 CNS gene sets enriched for schizophrenia as found by CEWAS displayed. The binary representation of the gene sets are projected onto 2D space using UMAP, and the colors indicate the level of gene set enrichment. (c) Top 25 CNS gene sets enriched for AD as found by CEWAS displayed in 2D UMAP space.

For neurodegenerative diseases, only CEWAS highlighted a CNS gene set related to abnormal spatial working memory for AD (Fig 5c), PD, and amyotrophic lateral sclerosis (ALS). Working memory deficit is a well recognized characteristic of AD^84^, and is also observed in PD^85^ and ALS^86^. Fittingly, genes found by CEWAS for AD, PD, and ALS (Table S6) are highly expressed in the parahippocampal gyrus, which relates to memory deficits seen in these conditions^87,88^. CEWAS also highlighted lateral medullary reticular group, cuneate nucleus, and gigantocellular nucleus for AD and PD, which aligns with how Lewy body accumulation in PD pathology starts in the reticular zone of the medulla oblongata^89^ and various brainstem nuclei are implicated in neurodegenerative diseases^90^. In fact, CEWAS highlighted a CNS gene set related to abnormal dopaminergic neuron morphology for AD, which aligns with how hippocampal activity is modulated by dopaminergic signals from brainstem, and dopaminergic cell death has been shown to correlate with memory deficits in AD mouse model^91^. Further, genes found by CEWAS for AD are highly expressed in the middle temporal gyrus and precuneus, which belong to the posterior default mode network, whose connectivity is disrupted in AD^92^. Lastly, CEWAS highlighted substantia nigra pars compacta for PD, which is the signature region of PD pathology where dopaminergic cell death occurs, resulting in the observed motor symptoms. Also, CEWAS highlighted a CNS gene set related to abnormal subventricular zone morphology for PD, which aligns with how neurogenesis in the subventricular zone is modulated by dopamine^51^.

## Discussion

We proposed CEWAS for integrating genotype, DNAm, and gene expression data to model the cascading effects of GWAS SNPs on DNAm, gene expression, and eventually phenotypes. We focused on this particular cascade of events since majority of GWAS SNPs lie in non-coding regions, hence their effects likely propagate from epigenome to transcriptome, as supported by previous mediation studies^15,93^. We built the CEWAS models using the largest brain tissue datasets comprising all three data types^20,21^. Higher detection rate was achieved by CEWAS compared to MetaXcan and EpiXcan, with high genomic correlation attained between well-powered, related GWAS. Also, while high correlations in z-scores between methods were observed, CEWAS identified numerous genes that are distinct from MetaXcan and EpiXcan, though some of these genes might be detectable with other methods not compared in this work. Stemming from large-scale GWAS, the distinct genes found by CEWAS are likely high-value targets, especially given that they are enriched for biologically relevant gene sets, and congruently, are highly expressed in disease-relevant brain regions.

Although CEWAS only captures epigenomic-mediated effects, this property could be useful for result interpretation. In particular, this property enables isolation of disease-relevant genes whose effects are regulated by a specific molecular mechanism, such as DNAm as in this study, which is the most widely examined molecular trait for gene regulation. As an example, *YPEL3* and *ZFYVE21* are significant for schizophrenia based on CEWAS but p>0.05 based on MetaXcan. CpGs selected by CEWAS include cg06985993 and cg08213375 for *YPEL3* and *ZFYVE21*, respectively, whose DNAm levels are associated with schizophrenia GWAS SNPs: rs3814877 and rs4900597^94^. Based on data from the same study, these SNPs are not associated with expression, hence suggesting their effects on schizophrenia might be mediated by DNAm. This result is supported by another study based on summary data-based Mendelian randomization^11^ that showed the effect of rs34813623 (LD = 0.84 with rs4900597) on schizophrenia is mediated by cg08213375 but not by expression. Though not pursued in this work, we can also isolate disease-relevant genes whose effects are regulated by histone acetylation, miRNA, and chromatin accessibility using the same approach. Hence, CEWAS could serve as a complement to existing methods that directly model genetic effects on gene expression in highlighting signals from the regulatory landscape of GWAS variants. Of note is that analogous to MetaXcan and other transcriptome wide association studies (TWAS), genes found by CEWAS might not be causal due to effects of LD between SNPs propagating to the predicted expression as well as shared GWAS variants across genes^95^.

CEWAS and EpiXcan might superficially appear to be similar, but the mechanisms being modeled and the actual information used are distinct. In CEWAS, we associate SNPs to CpGs and then CpGs to genes to explicitly model how certain GWAS SNPs affect gene regulation, which impacts gene expression and eventually the downstream phenotypes. In contrast, EpiXcan only weights SNPs based on epigenomic annotation during model learning. This distinction has two relevant implications. First, SNPs annotated to the same chromatin state are assigned the same sparsity weight in EpiXcan, whereas CEWAS estimates SNP-specific weights that reflect the association strengths between SNPs and CpGs. Second, a SNP selected by EpiXcan that resides on an annotated position might not necessarily be associated with epigenomic marks at or near that position, a property that CEWAS imposes. In fact, just using mQTL p-values as sparsity weights in EpiXcan, which captures the association strengths of SNPs on more distant CpGs, already increases detection rate over MetaXcan and EpiXcan but to a lesser extent than CEWAS (Fig S6). A plausible explanation for CEWAS’s higher detection rate compared to mQTL-weighted EpiXcan is that it enables more SNPs that are likely relevant to be modeled, since 543 samples are available per CpG for selecting relevant SNPs with elastic net and each gene is associated with 9.62 CpGs on average, whereas MetaXcan and EpiXcan have only 534 samples per gene for SNP selection (Fig S11).

Another method that is conceptually similar to CEWAS is MOSTWAS, but its focus is on incorporating *trans* genetic effects in predicting gene expression. Specifically, in addition to using *cis* SNPs, MOSTWAS uses *trans* SNPs associated with epigenomic mediators of each gene for expression model learning. Comparing the reported MOSTWAS z-scores (derived from models also built using ROSMAP data) against CEWAS, MetaXcan, and EpiXcan, correlations of 0.0691, 0.0949, and 0.1027 were found for AD2013, and correlations of 0.0159, 0.0061, and 0.0094 were found for MDD2018. In contrast, correlations between z-scores of CEWAS and MetaXcan/EpiXcan were 0.70 for AD2013 and 0.74 for MDD2018. Hence, incorporating *trans* genetic effects seem to highlight very different genes. We opted to focus on *cis* effects due to their higher replicability than *trans* effects^16^. Nonetheless, CEWAS can be expanded to incorporate longer distance relationships by including more distal epigenomic mediators of each gene in the expression prediction models. The respective merits of *cis* vs. *trans* is a huge topic that is beyond the scope of this study, but we acknowledge the importance of this topic, which deserves more in-depth future investigation.

Overall, we showed that CEWAS provides a simple yet effective way for integrating multi-omic data in estimating gene level z-scores from GWAS summary statistics. Numerous genes and gene sets highlighted by CEWAS point toward disease mechanisms that are promising but relatively less explored, e.g. mitochondria dysfunction in relation to metal ion binding and oxidative stress, neurogenesis impairment, and brain stem pathologies. All code and prediction models are made available (https://github.com/saramostafavi/CEWAS). This resource should prove useful for the research community to further investigate the regulatory landscape of GWAS variants in generating new research directions and finding new potential therapeutic targets for various brain-related disease and traits.

## Methods

### Genotype, DNA methylation, RNAseq, and GWAS data

For building the prediction models in CEWAS, we used genotype^17^, DNAm^18^ (Illumina 450K array), and RNAseq^19^ data from the ROSMAP study20,21. The genotype data were acquired from 2067 subjects. The DNAm and RNAseq data were derived from DLPFC tissue of 702 and 698 subjects, respectively. 543 subjects have both genotype and DNAm data, 485 subjects have both DNAm and RNAseq data, and 534 subjects have both genotype and RNAseq data. The data preprocessing pipelines are as previously described^15^ except we used the reference panel from the Haplotype Reference Consortium (HRC) to impute the genotype data, and ∼200 more subjects now have preprocessed RNAseq data available. We note that the top ten principal components (PC) were regressed out from the DNAm and RNAseq data as hidden confounders.

For replication, we used imputed genotype and preprocessed DLPFC-derived RNAseq data from the CMC^41^ study in combination with mQTLs from another large DLPFC tissue sample^42^, since DNAm data were not collected in the CMC study. 592 subjects have both genotype and RNAseq data in the CMC study, and the mQTLs were generated from 526 subjects. The DNAm data used in estimating the mQTLs were also acquired using Illumina 450K array.

For testing CEWAS, we used sixteen GWAS related to brain disease and traits^22–36^ (Table S1), namely schizophrenia (SCZ), bipolar disorder (BIP), AD, PD, ALS, depressive symptoms (DEP), major depressive disorder (MDD), wellbeing spectrum (wellbeing), insomnia, and intelligence (IQ). We refer to each GWAS by the disease/trait studied and the year at which the GWAS was performed, e.g. a schizophrenia GWAS performed in 2018 is referred to as SCZ2018.

### Cascading epigenomic analysis for GWAS

Motivated by the observation that GWAS SNPs are enriched in enhancers and open chromatin regions^2,96^, we propose CEWAS (Fig 1a) to analyze the cascading effects of genetics from epigenome to transcriptome and eventually to phenome. We first build a model to extract the genetic component of DNAm for each CpG *j*:

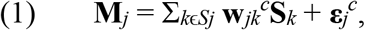

where **M**_*j*_ is a *n*×1 vector containing the DNAm levels of CpG *j*, **S**_*k*_ is a *n*×1 vector containing the dosage of SNP *k*, **w**_*jk*_^*c*^ is *k*^th^ element of a *l*_*j*_ ×1 model weight vector, **w**_*j*_^*c*^, to be estimated, and *S*_*j*_ is the set of *l*_*j*_ SNPs within ±50Kb from the CpG *j*. Following MetaXcan^9^, we estimated **w**_*jk*_^*c*^ using elastic net regression by applying GLMNET^38^ with its default settings, i.e. 10 fold cross-validation to set the sparsity parameter. To extract the epigenomic component of expression for each gene *i*, we modeled expression level in a similar manner:

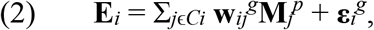

where **E**_*i*_ is a *n*×1 vector containing the expression level of gene *i* from *n* subjects, **M**_*j*_^*p*^ is a *n*×1 vector containing DNAm levels of CpG *j* predicted using (1), and **w**_*ij*_^*g*^ is the *j*^th^ element of a *m*_*i*_×1 model weight vector, **w**_*i*_^*g*^, to be estimated with elastic net. *C*_*i*_ is the set of *m*_i_ CpGs within ±500Kb from the TSS of gene *i*. Note that window sizes were chosen so that SNPs selected by CEWAS would mostly lie within a typical 1Mb window from TSS as used in MetaXcan^9^ and EpiXcan^13^. Also, to estimate the R^2^ for each gene, we applied (1) and (2) sequentially and computed the square of correlation between the predicted and measured expression levels^97^. Only genes with R^2^ > 0.01 were retained for analysis.

Given **w**_*ij*_^*g*^ and **w**_*jk*_^*c*^, genetically-driven epigenomic effects at gene level can be estimated in a manner analogous to sequentially applying MetaXcan:

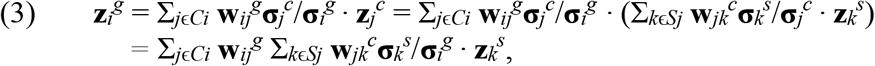

where **z**_*i*_^*g*^ is the z-score at gene level for gene *i*, **z**_*j*_^*c*^ is the z-score at CpG level for CpG *j*, **z**_*k*_^*s*^ is the z-score at SNP for SNP *k*. **σ**_*i*_^*g*^ and **σ**_*k*_^*s*^ are the variance of gene *i* and SNP *k*, respectively. We highlight that since only genetically-driven epigenomic effects are retained by (1), we must estimate **σ**_*i*_^*g*^ based on the genetic component of DNAm. For this, we set **σ**_*i*_^*g*^ to **w**_*i*_^*g*T^*cov*(**M**^*p*^)**w**_*i*_^*g*^, where **M**^*p*^ is a *n*×*m*^i^ matrix containing DNAm levels predicted using (1). As shown in the Fig 2, estimating **σ**_*i*_^*g*^ with only the genetic component of DNAm is critical for **z**_*i*_^*g*^ to be calibrated.

### Gene detection and distinct signal estimation

We applied (3) to sixteen well-powered, brain-related GWAS^22–36^. Genes were declared significant at an *α* of 0.05 with Bonferroni correction for the number of tested genes. Since some genes might have non-zero weights assigned to the same SNPs, we also estimated the number of distinct signals within the significant genes and within the tested genes to provide another estimate of detection rate (Fig S1). To estimate the number of distinct signals, we used an approach that combines permutation test with principal component analysis (PCA)^98^. Specifically, we first applied PCA to the predicted expression of queried genes (significant or tested) for each method, and recorded the percentage variance explained by each PC. We then permuted the predicted expression matrix along both rows and columns a thousand times to generate the null distribution of the percentage variance for each PC as used for estimating p-values. PCs with p < 0.05 were declared significant.

We note that due to difficulties in genotyping the MHC region, genes (defined as ±500Kb from TSS) that overlap with the MHC region were excluded from analysis. Also, although the ROSMAP subjects are of European descent, the reference allele for some SNPs could still be different from those of GWAS. We thus accounted for allele flips by inverting the signs of the GWAS z-scores, and removed all ambiguous SNPs, i.e. cases where A1 and A2 are complementary, e.g. A1=A, A2=T. For comparison, we applied MetaXcan with expression prediction models, **w**_*ik*_^*g*^, built using the same SNP sets as CEWAS as well as SNPs within ±500Kb from TSS of each gene *i*. The former ensures the same SNPs as CEWAS are considered to test the effect of incorporating DNAm, while the latter follows conventions in the literature^99^. For comparison against EpiXcan, similar expression prediction models were built using elastic net except the sparse penalty was weighted based on epigenomic annotation of the SNPs^13^. Only genes with R^2^ > 0.01 were kept.

### z-score calibration test

For the detected genes to be trustworthy, we need to ensure that **z**_*i*_^*g*^ generated by (3) is calibrated. In particular, applying (3) to null **z**_*k*_^*s*^ should output null **z**_*i*_^*g*^. To test whether this criterion is met, special care is needed in simulating null **z**_*k*_^*s*^. Specifically, in addition to requiring **z**_*k*_^*s*^ of each SNP *k* to follow *N*(0,1), the LD structure of all SNPs involved in (3) must be accounted for. Otherwise, the correlations between CpGs would not be properly modeled. To satisfy these two conditions, for each gene *i*, we drew 10000 sets of **z**_*k*_^*s*^ from *N*(**0**,**R**_*i*_), where **R**_*i*_ is the correlation between all SNPs in {*S*_*j*_} for *j* ϵ *C*_*i*_. Using correlation, as opposed to covariance, ensures the standard deviation of each **z**_*k*_^*s*^ is 1. For generating **R**_*i*_, we used the ROSMAP imputed genotype data. To evaluate the resulting **z**_*i*_^*g*^, we checked if each set of 10000 **z**_*i*_^*g*^’s of each gene *i* follows *N*(0,1). In practice, the LD structure of GWAS genotype data might not match that of ROSMAP genotype data. To test the effect of LD mismatch, we further used **R**_*i*_ estimated from the 1000 Genome phase 3 genotype data (European population) to generate **z**_*k*_^*s*^.

### Replication

We used the CMC data^41^ and mQTLs from another large DLPFC tissue sample^42^ for replication. To align the allele across the CMC data, the mQTLs, and the GWAS, we matched the allele of all these datasets to that of the 1000 Genomes panel^100^. All regression coefficients, **β**_*ik*_^*c*^, associated with each CpG *j* in the mQTL set were used as **w**_*j*_^*c*^ in (3). To estimate **w**_*i*_^*g*^, we first multiplied **β**_*jk*_^*c*^ to the corresponding SNPs in the CMC data (with allele matched) to generate predicted DNAm levels. We then applied elastic net regression to these predicted DNAm levels and the CMC gene expression data to generate **w**_*i*_^*g*^. **σ**_*k*_^*s*^ was estimated using the 1000 Genomes data, and **σ**_*i*_^*g*^ was estimated using the predicted DNAm levels. We applied (3) with these model parameters to the same sixteen GWAS^22–36^, and used area under the receiver operating characteristic curve (AUC) as the replication metric. To estimate AUC, we ranked the genes of each GWAS based on p-values estimated with the ROSMAP models, took the top 1% of genes as the reference, and computed true positive rate and false positive rate by applying a range of thresholds from 0 to 1 on p-values derived from the CMC models. We took the top 1% of genes as reference since less than a handful of genes are significant for some GWAS, which might be too few for robust AUC estimation. We further assessed statistical significance by estimating AUC on 10000 sets of permuted p-values and confirmed that chance level AUC is 0.5±0.02. For comparison, we evaluated the replication of MetaXcan^99^ and EpiXcan^13^ derived from ROSMAP vs. CMC data, with the latter being provided by the respective authors. ROSMAP models built with SNPs within ±500Kb from TSS were used to match the way the CMC models were built. We also examined the correlations between **z**_*i*_^*g*^’s across related GWAS as another form of replication assessment. Note that well being spectrum is partly estimated based on depressive symptoms^35^, hence **z**_*i*_^*g*^’s of wellbeing2019 are expected to be highly correlated with DEP2019, MDD2019, and MDD2018.

We note that the lack of another large brain tissue dataset with genotype, DNAm, and gene expression data complicates replication. On the model learning side, we must first align the allele of the CMC genotype data and the mQTLs to the same reference for generating (predicted) DNAm data. This alignment process includes flipping signs of mQTLs and flipping dosage coding of CMC genotype data whenever allele mismatches occur, which is prone to errors especially for SNPs with allele frequency close to 0.5. Also, mQTLs of ambiguous SNPs need to be dropped, i.e. cases where A1 and A2 are complementary, e.g. A1=A, A2=T. On the z-score estimation side, we must align the allele of the GWAS SNPs to the same reference used for model learning, which is again prone to sign flip errors and variable mismatches.

### Gene set enrichment

To infer the biological processes captured by CEWAS, we examined gene set enrichment by adapting the contrastive analysis proposed in MAGMA^70^. Specifically, we used generalized least square to model the gene level z-scores:

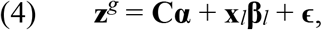

where **z**^*g*^ is a *q*×1 vector containing the gene level z-scores, **C** is a *q*×*d* matrix containing *d* confounds, **x**_*l*_ is a *q*×1 binary vector with 1 indicating genes that belong to gene set *l*, **ϵ** ∼ *N*(0,**Σ**), and **Σ** is a *q*×*q* matrix to account for correlation between genes. We used **C** to account for the number of SNPs selected by CEWAS for each given gene (i.e. Σ_*jk*_ (**w**_*ij*_**w**_*jk*_ > 0)), and we applied oracle approximating shrinkage^101^ to the gene expression data to estimate a well-conditioned **Σ** that is closest to the unknown ground truth **Σ** in the least square sense. Having a well-conditioned estimate of **Σ** is critical since **Σ**^-1^ is involved in the estimation of **β**_*l*_ and se(**β**_*l*_). CNS gene sets^22^ as well as GO and canonical gene sets (MSigDB v7.1^71^) having 10 to 200 genes^102^ (that intersect with the tested genes of each method) were examined. Significance was declared at an α of 0.05 with FDR correction for the number of gene sets in the CNS set and in each MSigDB collection.

### Brain region association

We also examined association with brain regions by applying (4) to the human brain microarray data from Allen Institute^72^. Given gene-by-region expression matrices derived from six post-mortem brains, we first averaged the expression values corresponding to the same gene-region pairs for each subject. We then applied Student’s t-test to the average expression values of each unique gene-region pairs across subjects. The vector of t-values for each region was taken as **x**_*l*_ in (4) to estimate brain region association. Only genes tested by CEWAS and available in the Allen Institute data were used in (4). The same analysis was performed with MetaXcan and EpiXcan z-scores. Significance was declared at an α of 0.05 with FDR correction for the number of regions.

### Software availability

All results are generated using in-house MATLAB scripts. To increase portability, we have built a python software that interfaces with the widely-used MetaXcan software, and assembled all CEWAS models and parameters into the required format. The python software is available on our GitHub page: https://github.com/saramostafavi/CEWAS. CEWAS can be executed with a single command using this software and takes between 4-10 min to run for a typical GWAS dataset on a 4 core machine. Installation details with a sample test case are provided on the GitHub page.

## Accession Codes

Genotype, RNA-seq, and DNAm data for ROSMAP samples are available from the Synapse AMP-AD Data Portal https://www.synapse.org/#!Synapse:syn2580853/discussion/default as well as RADC Research Resource Sharing Hub at www.radc.rush.edu.

## Acknowledgements

We thank the participants of ROS and MAP for their essential contributions and gift to this project. This work has been supported by many different NIH grants: P30AG10161, U01 AG046152, U01AG61356, R01 AG057911, R01 AG061798, RC2AG036547. The funders had no role in study design, data collection and analysis, decision to publish, or preparation of the manuscript.

## Contributions

Study design: SM, BN. Sample collection: DAB. Data generation and quality control analyses: BN, CG, PLD, ST. Analyses: BN, SM, CW, NHK, FF. Interpretation of results and critical review of the manuscript: BN, CG, ST, PLD, DAB.

## Supplementary Tables

**Table S1. Sixteen brain-related GWAS analyzed**

**Table S2. GWAS and detection summary**

**Table S3. Replication AUC**

**Table S4. Gene detection statistics**

**Table S5. Gene set enrichment**

**Table S6. Brain region enrichment**

## Supplementary Figure Legends

**Fig S1. Detection rate based on distinct signals**. Ratio of the number of significant PCs extracted from the significant genes over the number of significant PCs extracted from the tested genes displayed. The number of significant PCs was used as an estimate of the number of distinct signals within a set of genes (see Methods). This analysis accounts for how some SNPs are shared between models of spatially proximal genes. The overall trend of CEWAS attaining higher detection rate than MetaXcan and EpiXcan remains with this analysis.

**Fig S2. R**^**2**^ **of gene models**. (a) R^2^ estimated by correlating the predicted and measured expression levels in ROSMAP. CEWAS attained lower R^2^ as expected, since CEWAS is designed to extract a specific component of gene expression, namely the epigenomic component that is driven by genetic effects. In contrast, MetaXcan and EpiXcan are optimized for predicting gene expression. (b) R^2^ estimated by applying ROSMAP models to CMC genotype data and correlating the predicted expression levels with that measured in CMC. R^2^ appears similar across methods, but CEWAS actually attained slightly lower R^2^ if we zoom into the results (see Fig S3).

**Fig S3. Correlation between predicted and measured expression levels on CMC data**. Models trained with ROSMAP data were applied to CMC genotype data to predict gene expression. Correlation attained by CEWAS is slightly lower than MetaXcan and EpiXcan.

**Fig S4. Stability analysis of partitioned heritability**. Bars show the log p-value of enrichment of mQTLs and eQTLs assessed by LDSC for each GWAS. (a) The enrichment for mQTLs remains higher than for eQTLs with the window size of eQTLs matched to that of mQTLs (p=0.0084, Wilcoxon sign rank test across GWAS). (b) The same trend holds but to a lesser extent when we matched the number of mQTL SNPs to the number of eQTL SNPs by taking mQTL SNPs that are closer to CpGs (p=0.1122, Wilcoxon sign rank test across GWAS; p=0.0269, paired t-test).

**Fig S5. Replication of mQTLs vs. eQTLs**. Using mQTLs in Jaffe et al.^35^ as the reference, we first found the top mQTL SNP for each CpG, and ordered the resulting top mQTLs based on p-values. Taking *q*% of the top mQTLs as “ground truth”, *q* = 5% to 50%, we computed AUC on mQTL p-values derived from the ROSMAP data. Since mQTLs from Jaffe et al. were estimated with a 20Kb window, we restricted the ROSMAP mQTLs to those within the same window. The same procedure was applied to estimate AUC for eQTLs, with eQTLs from CMC as the reference. Since eQTLs from CMC were estimated with a 1Mb window, we first examined ROSMAP eQTLs within a 1Mb window, but also examined CMC and ROSMAP eQTLs restricted to the same window size as the mQTLs, i.e. 20Kb. To estimate the null, we permuted the top eQTLs/mQTLs and repeated the procedure.

**Fig S6. Gene detection rate with DNAm mediated effects on GWAS phenotypes**. We assessed DNAm mediated effects on GWAS phenotypes in four ways. First, we examined results from models used in the first stage of CEWAS, i.e. MetaXcan models built with each CpG taken as the response and SNPs within ±50Kb from that CpG as predictors. Detection rate of these DNAm MetaXcan models, defined as the number of significant CpGs among tested CpGs, is significantly lower than CEWAS (p=0.0004), MetaXcan (p=0.0013), and EpiXcan (p=0.0006) based on Wilcoxon sign rank test across GWAS. Second, we assessed CpG-to-gene mapping that does not use expression data, by taking the DNAm MetaXcan p-value of the closest CpG of each gene as the p-value of that gene. The gene level detection rate is significantly lower than CEWAS (p=0.0004), MetaXcan (p=0.0003), and EpiXcan (p=0.0004). Third, we took the DNAm MetaXcan p-value of the CpG with the largest R^2^ in terms of DNAm prediction among CpGs within ±500Kb from each gene as the p-value of that gene. The detection rate is lower than CEWAS (p=0.1331) and EpiXcan (p=0.3808), and higher than MetaXcan (p=0.4235). Fourth, to contrast using epigenomic annotation against explicitly modeling associations between SNPs and CpGs, we modified EpiXcan by weighting the sparse penalty using mQTL p-values. We first associated each CpG to SNPs that are within ±50Kb. We then found the smallest p-value, *pk*, across CpGs for each SNP *k*, and used 10^10^*pk*+0.5 (capped at 1) as the weight for its sparse penalty. Multiplying by 10^10^ accounts for the number of SNP-CpG pairs tested, and +0.5 penalizes SNPs with the strongest mQTL effects by half the amount as SNPs with weak/no mQTL effects. The detection rate of mQTL-weighted EpiXcan is significantly higher than both MetaXcan (p=0.00006) and EpiXcan (p=0.00006), and lower than CEWAS on average (p=0.3881).

**Fig S7. Genomic correlation between schizophrenia and IQ**. Gene level z-scores estimated by CEWAS for schizophrenia vs. IQ displayed. A genomic correlation of -0.15 was observed. The genes that were detected by CEWAS across schizophrenia, bipolar disorder, depression, and IQ are highlighted in bold. Observing opposite signs in z-scores for the highlighted genes matches how risk alleles in these loci were shown to correlate with lower cognitive test scores^37^.

**Fig S8. Number of distinct signals among differential genes**. Number of distinct signals (i.e. number of significant PCs, see Methods) among differential genes (darker shade) and significant genes (lighter shade) detected by each method. Only genes tested in all three methods were considered in extracting differential genes exclusively found by one method but not the other two. The overall trend of CEWAS finding more distinct signals than MetaXcan and EpiXcan remains.

**Fig S9. Comparison with spatially mapping GWAS SNPs**. (a) The distance between TSS of differential genes exclusively found by CEWAS and their closest GWAS hits summarized. Each bar corresponds to the percentage of differential genes lying in a range of distance, e.g. a bar between 100 and 1000 corresponds to the percentage of differential genes with distance between 100 and 1000 base pairs. More than half of the differential genes are >100Kb away from any GWAS hits. (b) The percentage of differential genes exclusively found by CEWAS but missed by spatially mapping GWAS SNPs to their closest genes shown. Note that no differential genes were found for ALS2018, hence why the percentage is zero.

**Fig S10. Correlation of gene set enrichment scores between CEWAS and contrasted methods**. The average correlation over GWAS for each gene set category displayed. The observed correlation suggests moderate similarity in enriched gene sets between CEWAS and the contrasted methods, which we confirmed by manual inspection.

**Fig S11. Number of selected SNPs**. (a) Number of SNPs selected per CpG by elastic net. (b) Number of SNP selected per gene by elastic net. CEWAS tends to have more SNPs selected since 543 samples are available per CpG for selecting relevant SNPs with elastic net and each gene is associated with 9.62 CpGs on average, whereas MetaXcan and EpiXcan have only 534 samples per gene for SNP selection.

## Reference

1. Hindorff, L. A. et al. Potential etiologic and functional implications of genome-wide association loci for human diseases and traits. Proc. Natl. Acad. Sci. 106, 9362–9367 (2009).

2. Ernst, J. et al. Mapping and analysis of chromatin state dynamics in nine human cell types. Nature 473, 43 (2011).

3. Nica, A. C. et al. Candidate causal regulatory effects by integration of expression QTLs with complex trait genetic associations. PLoS Genet. 6, e1000895 (2010).

4. Giambartolomei, C. et al. Bayesian test for colocalisation between pairs of genetic association studies using summary statistics. PLoS Genet. 10, e1004383 (2014).

5. Gleason, K. J., Yang, F., Pierce, B. L., He, X. & Chen, L. S. Primo: integration of multiple GWAS and omics QTL summary statistics for elucidation of molecular mechanisms of trait-associated SNPs and detection of pleiotropy in complex traits. Genome Biol. 2020 211 21, 1–24 (2020).

6. Giambartolomei, C. et al. A Bayesian framework for multiple trait colocalization from summary association statistics. Bioinformatics 34, 2538–2545 (2018).

7. Foley, C. N. et al. A fast and efficient colocalization algorithm for identifying shared genetic risk factors across multiple traits. Nat. Commun. 2021 121 12, 1–18 (2021).

8. Gusev, A. et al. Integrative approaches for large-scale transcriptome-wide association studies. Nat. Genet. 48, 245 (2016).

9. Barbeira, A. N. et al. Exploring the phenotypic consequences of tissue specific gene expression variation inferred from GWAS summary statistics. Nat. Commun. 9, 1–20 (2018).

10. Hannon, E. et al. Leveraging DNA-methylation quantitative-trait loci to characterize the relationship between Methylomic variation, gene expression, and complex traits. Am. J. Hum. Genet. 103, 654–665 (2018).

11. Hannon, E., Weedon, M., Bray, N., O’Donovan, M. & Mill, J. Pleiotropic effects of trait-associated genetic variation on DNA methylation: utility for refining GWAS loci. Am. J. Hum. Genet. 100, 954–959 (2017).

12. Wu, Y. et al. Integrative analysis of omics summary data reveals putative mechanisms underlying complex traits. Nat. Commun. 9, 918 (2018).

13. Zhang, W. et al. Integrative transcriptome imputation reveals tissue-specific and shared biological mechanisms mediating susceptibility to complex traits. Nat. Commun. 10, (2019).

14. Bhattacharya, A., Li, Y. & Love, M. I. MOSTWAS: Multi-Omic Strategies for Transcriptome-Wide Association Studies. PLOS Genet. 17, e1009398 (2021).

15. Ng, B. et al. An xQTL map integrates the genetic architecture of the human brain’s transcriptome and epigenome. Nat. Neurosci. 20, 1418–1426 (2017).

16. Joehanes, R. et al. Integrated genome-wide analysis of expression quantitative trait loci aids interpretation of genomic association studies. Genome Biol. 2017 181 18, 1–24 (2017).

17. Farfel, J. M. et al. Relation of genomic variants for Alzheimer disease dementia to common neuropathologies. Neurology 87, 489–496 (2016).

18. De Jager, P. L. et al. Alzheimer’s disease: early alterations in brain DNA methylation at ANK1, BIN1, RHBDF2 and other loci. Nat. Neurosci. 17, 1156–1163 (2014).

19. Mostafavi, S. et al. A molecular network of the aging human brain provides insights into the pathology and cognitive decline of Alzheimer’s disease. Nat. Neurosci. 21, 811–819 (2018).

20. Bennett, D. A. et al. Religious orders study and rush memory and aging project. J. Alzheimer’s Dis. 1–28 (2018).

21. De Jager, P. L. et al. Data descriptor: A multi-omic atlas of the human frontal cortex for aging and Alzheimer’s disease research. Sci. Data 5, (2018).

22. Pardiñas, A. F. et al. Common schizophrenia alleles are enriched in mutation-intolerant genes and in regions under strong background selection. Nat. Genet. 50, 381 (2018).

23. Biological insights from 108 schizophrenia-associated genetic loci. Nature 511, 421–427 (2014).

24. Wray, N. R. et al. Genome-wide association analyses identify 44 risk variants and refine the genetic architecture of major depression. Nat. Genet. 50, 668 (2018).

25. Jansen, P. R. et al. Genome-wide analysis of insomnia in 1,331,010 individuals identifies new risk loci and functional pathways. Nat. Genet. 51, 394–403 (2019).

26. Hammerschlag, A. R. et al. Genome-wide association analysis of insomnia complaints identifies risk genes and genetic overlap with psychiatric and metabolic traits. Nat. Genet. 49, 1584–1592 (2017).

27. Savage, J. E. et al. Genome-wide association meta-analysis in 269,867 individuals identifies new genetic and functional links to intelligence. Nat. Genet. 50, 912–919 (2018).

28. Sniekers, S. et al. Genome-wide association meta-A nalysis of 78,308 individuals identifies new loci and genes influencing human intelligence. Nat. Genet. 49, 1107–1112 (2017).

29. Stahl, E. A. et al. Genome-wide association study identifies 30 loci associated with bipolar disorder. Nat. Genet. 51, 793–803 (2019).

30. Jansen, I. E. et al. Genome-wide meta-analysis identifies new loci and functional pathways influencing Alzheimer’s disease risk. (2019).

31. Marioni, R. E. et al. GWAS on family history of Alzheimer’s disease. Transl. Psychiatry 8, (2018).

32. Lambert, J. C. et al. Meta-analysis of 74,046 individuals identifies 11 new susceptibility loci for Alzheimer’s disease. Nat Genet 45, 1452–1458 (2013).

33. Nalls, M. A. et al. Expanding Parkinson’s disease genetics: novel risk loci, genomic context, causal insights and heritable risk. bioRxiv 388165 (2019) doi:10.1101/388165.

34. Nicolas, A. et al. Genome-wide Analyses Identify KIF5A as a Novel ALS Gene. Neuron 97, 1268–1283.e6 (2018).

35. Baselmans, B. M. L. et al. Multivariate genome-wide analyses of the well-being spectrum. Nat. Genet. 51, 445–451 (2019).

36. Howard, D. M. et al. Genome-wide meta-analysis of depression identifies 102 independent variants and highlights the importance of the prefrontal brain regions. Nat. Neurosci. 22, 343–352 (2019).

37. Battle, A. et al. Characterizing the genetic basis of transcriptome diversity through RNA-sequencing of 922 individuals. Genome Res. 24, 14–24 (2014).

38. Friedman, J. & Tibshirani, R. Regularization paths for generalized linear models via coordinate descent. J. Stat. Softw. 33, 1–22 (2010).

39. Finucane, H. K. et al. Partitioning heritability by functional annotation using genome-wide association summary statistics. Nat Genet 47, 1228–1235 (2015).

40. Hormozdiari, F. et al. Leveraging molecular quantitative trait loci to understand the genetic architecture of diseases and complex traits. Nat. Genet. 2018 507 50, 1041–1047 (2018).

41. Fromer, M. et al. Gene expression elucidates functional impact of polygenic risk for schizophrenia. Nat Neurosci 19, 1442–1453 (2016).

42. Jaffe, A. E. et al. Mapping DNA methylation across development, genotype and schizophrenia in the human frontal cortex. Nat. Neurosci. 19, 40 (2016).

43. N, M. et al. Integrating Gene Expression with Summary Association Statistics to Identify Genes Associated with 30 Complex Traits. Am. J. Hum. Genet. 100, 473–487 (2017).

44. Ruderfer, D. M. et al. Genomic Dissection of Bipolar Disorder and Schizophrenia, Including 28 Subphenotypes. Cell 173, 1705–1715.e16 (2018).

45. Yang, Z. et al. The genome-wide risk alleles for psychiatric disorders at 3p21.1 show convergent effects on mRNA expression, cognitive function, and mushroom dendritic spine. Mol. Psychiatry 25, 48–66 (2020).

46. Smith, D. G. et al. Melanin-concentrating hormone-1 receptor modulates neuroendocrine, behavioral, and corticolimbic neurochemical stress responses in mice. Neuropsychopharmacology 31, 1135–1145 (2006).

47. Guerreiro, R. et al. Genome-wide analysis of genetic correlation in dementia with Lewy bodies, Parkinson’s and Alzheimer’s diseases. Neurobiol. Aging 38, 214.e7-214.e10 (2016).

48. Peng, G. et al. Genome-wide transcriptome profiling of homologous recombination DNA repair. Nat. Commun. 5, 1–11 (2014).

49. Urrutia, R. KRAB-containing zinc-finger repressor proteins. Genome Biology vol. 4 231 (2003).

50. Moreno-Jiménez, E. P. et al. Adult hippocampal neurogenesis is abundant in neurologically healthy subjects and drops sharply in patients with Alzheimer’s disease. Nat. Med. 25, 554–560 (2019).

51. Höglinger, G. U. et al. Dopamine depletion impairs precursor cell proliferation in Parkinson disease. Nat. Neurosci. 7, 726–735 (2004).

52. Smith, C. L. & Eppig, J. T. The mammalian phenotype ontology: Enabling robust annotation and comparative analysis. Wiley Interdiscip. Rev. Syst. Biol. Med. 1, 390–399 (2009).

53. Murphy, M. P. How mitochondria produce reactive oxygen species. Biochemical Journal vol. 417 1–13 (2009).

54. Bitanihirwe, B. K. Y. & Woo, T.-U. W. Oxidative Stress in Schizophrenia: An Integrated Approach. Neurosci. Biobehav. Rev. 35, 878 (2011).

55. M, B. et al. Pathways underlying neuroprogression in bipolar disorder: focus on inflammation, oxidative stress and neurotrophic factors. Neurosci. Biobehav. Rev. 35, 804–817 (2011).

56. Lammich, S. et al. Constitutive and regulated α-secretase cleavage of Alzheimer’s amyloid precursor protein by a disintegrin metalloprotease. Proc. Natl. Acad. Sci. U. S. A. 96, 3922–3927 (1999).

57. Gregg, S. Q., Robinson, A. R. & Niedernhofer, L. J. Physiological consequences of defects in ERCC1-XPF DNA repair endonuclease. DNA Repair vol. 10 781–791 (2011).

58. Lam, Y. A. et al. Inhibition of the ubiquitin-proteasome system in Alzheimer’s disease. Proc. Natl. Acad. Sci. U. S. A. 97, 9902–9906 (2000).

59. McEwan, D. G. et al. PLEKHM1 regulates autophagosome-lysosome fusion through HOPS complex and LC3/GABARAP proteins. Mol. Cell 57, 39–54 (2015).

60. Dinter, E. et al. Rab7 induces clearance of α-synuclein aggregates. J. Neurochem. 138, 758–774 (2016).

61. Ghezzi, D. et al. Mutations in TTC19 cause mitochondrial complex III deficiency and neurological impairment in humans and flies. Nature Genetics vol. 43 259–263 (2011).

62. Zhou, C., Huang, Y. & Przedborski, S. Oxidative stress in Parkinson’s disease: A mechanism of pathogenic and therapeutic significance. in Annals of the New York Academy of Sciences vol. 1147 93–104 (Blackwell Publishing Inc., 2008).

63. Dias, C. et al. β-catenin mediates stress resilience through Dicer1/microRNA regulation. Nature 516, S1–S5 (2014).

64. Gorinski, N. et al. Attenuated palmitoylation of serotonin receptor 5-HT1A affects receptor function and contributes to depression-like behaviors. Nat. Commun. 10, 1–14 (2019).

65. Park, C. Y. et al. Genome-wide landscape of RNA-binding protein target site dysregulation reveals a major impact on psychiatric disorder risk. Nat. Genet. 2021 532 53, 166–173 (2021).

66. MahmoudianDehkordi, S. et al. Altered Bile Acid Profile Associates with Cognitive Impairment in Alzheimer’s Disease – An Emerging Role for Gut Microbiome. Alzheimers. Dement. 15, 76 (2019).

67. Baloni, P. et al. Metabolic Network Analysis Reveals Altered Bile Acid Synthesis and Metabolism in Alzheimer’s Disease. Cell Reports Med. 1, (2020).

68. Mukai, J. et al. Recapitulation and Reversal of Schizophrenia-Related Phenotypes in Setd1a-Deficient Mice. Neuron 104, 471–487.e12 (2019).

69. Nagahama, K. et al. Setd1a Insufficiency in Mice Attenuates Excitatory Synaptic Function and Recapitulates Schizophrenia-Related Behavioral Abnormalities. Cell Rep. 32, 108126 (2020).

70. de Leeuw, C. A., Mooij, J. M., Heskes, T. & Posthuma, D. MAGMA: Generalized Gene-Set Analysis of GWAS Data. PLOS Comput. Biol. 11, e1004219 (2015).

71. Liberzon, A. et al. The Molecular Signatures Database Hallmark Gene Set Collection. Cell Syst. 1, 417–425 (2015).

72. Hawrylycz, M. J. et al. An anatomically comprehensive atlas of the adult human brain transcriptome. Nature 489, 391–399 (2012).

73. Cotter, D. R., Pariante, C. M. & Everall, I. P. Glial cell abnormalities in major psychiatric disorders: The evidence and implications. Brain Research Bulletin vol. 55 585–595 (2001).

74. Braff, D. et al. Prestimulus Effects on Human Startle Reflex in Normals and Schizophrenics. Psychophysiology 15, 339–343 (1978).

75. Perry, W., Minassian, A., Feifel, D. & Braff, D. L. Sensorimotor gating deficits in bipolar disorder patients with acute psychotic mania. Biol. Psychiatry 50, 418–424 (2001).

76. Perry, W., Minassian, A. & Feifel, D. Prepulse inhibition in patients with non-psychotic major depressive disorder. J. Affect. Disord. 81, 179–184 (2004).

77. Yun, S., Reynolds, R. P., Masiulis, I. & Eisch, A. J. Re-evaluating the link between neuropsychiatric disorders and dysregulated adult neurogenesis. Nature Medicine vol. 22 1239–1247 (2016).

78. Lee, Y. & Davis, M. Role of the hippocampus, the bed nucleus of the stria terminalis, and the amygdala in the excitatory effect of corticotropin-releasing hormone on the acoustic startle reflex. J. Neurosci. 17, 6434–6446 (1997).

79. Gross, C. G. Neurogenesis in the adult brain: Death of a dogma. Nat. Rev. Neurosci. 1, 67–73 (2000).

80. Lucki, I. The spectrum of behaviors influenced by serotonin. Biol. Psychiatry 44, 151–162 (1998).

81. Steullet, P. et al. The thalamic reticular nucleus in schizophrenia and bipolar disorder: role of parvalbumin-expressing neuron networks and oxidative stress. Mol. Psychiatry 23, 2057–2065 (2018).

82. Howes, O. D. & Kapur, S. The Dopamine Hypothesis of Schizophrenia: Version III-The Final Common Pathway. doi:10.1093/schbul/sbp006.

83. Dunlop, B. W. & Nemeroff, C. B. The role of dopamine in the pathophysiology of depression. Archives of General Psychiatry vol. 64 327–337 (2007).

84. Baddeley, A. D., Bressi, S., Della Sala, S., Logie, R. & Spinnler, H. THE DECLINE OF WORKING MEMORY IN ALZHEIMER’S DISEASE A LONGITUDINAL STUDY. Brain vol. 114 https://academic.oup.com/brain/article-abstract/114/6/2521/371418 (1991).

85. Cools, R. Dopaminergic modulation of cognitive function-implications for L-DOPA treatment in Parkinson’s disease. Neuroscience and Biobehavioral Reviews vol. 30 1–23 (2006).

86. Goldstein, L. H. & Abrahams, S. Changes in cognition and behaviour in amyotrophic lateral sclerosis: Nature of impairment and implications for assessment. The Lancet Neurology vol. 12 368–380 (2013).

87. Takeda, T., Uchihara, T., Arai, N., Mizutani, T. & Iwata, M. Progression of hippocampal degeneration in amyotrophic lateral sclerosis with or without memory impairment: Distinction from Alzheimer disease. Acta Neuropathol. 117, 35–44 (2009).

88. Lè Ne Hall, H. et al. Hippocampal Lewy pathology and cholinergic dysfunction are associated with dementia in Parkinson’s disease. A J. Neurol. doi:10.1093/brain/awu193.

89. Seidel, K. et al. The brainstem pathologies of Parkinson’s disease and dementia with lewy bodies. Brain Pathol. 25, 121–135 (2015).

90. Grinberg, L. T., Rueb, U. & Heinsen, H. Brainstem: Neglected locus in neurodegenerative diseases. Front. Neurol. JUL, (2011).

91. Nobili, A. et al. Dopamine neuronal loss contributes to memory and reward dysfunction in a model of Alzheimer’s disease. Nat. Commun. 8, 1–14 (2017).

92. Greicius, M. D., Srivastava, G., Reiss, A. L. & Menon, V. Default-mode network activity distinguishes Alzheimer’s disease from healthy aging: evidence from functional MRI. Proc. Natl. Acad. Sci. U. S. A. 101, 4637–4642 (2004).

93. Hemani, G., Tilling, K. & Smith, G. D. Orienting the causal relationship between imprecisely measured traits using GWAS summary data. PLOS Genet. 13, e1007081 (2017).

94. Lin, D. et al. Characterization of cross-tissue genetic-epigenetic effects and their patterns in schizophrenia. Genome Med. 2018 101 10, 1–12 (2018).

95. Wainberg, M. et al. Opportunities and challenges for transcriptome-wide association studies. Nat. Genet. 51, 592 (2019).

96. Schaub, M. A., Boyle, A. P., Kundaje, A., Batzoglou, S. & Snyder, M. Linking disease associations with regulatory information in the human genome. Genome Res. 22, 1748–1759 (2012).

97. Gamazon, E. R. et al. A gene-based association method for mapping traits using reference transcriptome data. Nat. Genet. 47, 1091 (2015).

98. Mao, W., Zaslavsky, E., Hartmann, B. M., Sealfon, S. C. & Chikina, M. Pathway-level information extractor (PLIER) for gene expression data. Nat. Methods 16, 607–610 (2019).

99. Barbeira, A. N. et al. Exploring the phenotypic consequences of tissue specific gene expression variation inferred from GWAS summary statistics. Nat. Commun. 9, 1825 (2018).

100. Abecasis, G. R. et al. An integrated map of genetic variation from 1,092 human genomes. Nature 491, 56–65 (2012).

101. Chen, Y., Wiesel, A., Eldar, Y. C. & Hero, A. O. Shrinkage algorithms for MMSE covariance estimation. IEEE Trans. Sig. Proc. 58, 5016–5029 (2010).

102. Reimand, J. et al. Pathway enrichment analysis and visualization of omics data using g:Profiler, GSEA, Cytoscape and EnrichmentMap. Nat. Protoc. 14, 482–517 (2019).

