## Supplementary Figures for "Cascading Epigenomic Analysis for Identifying Disease Genes from the Regulatory Landscape of GWAS Variants"

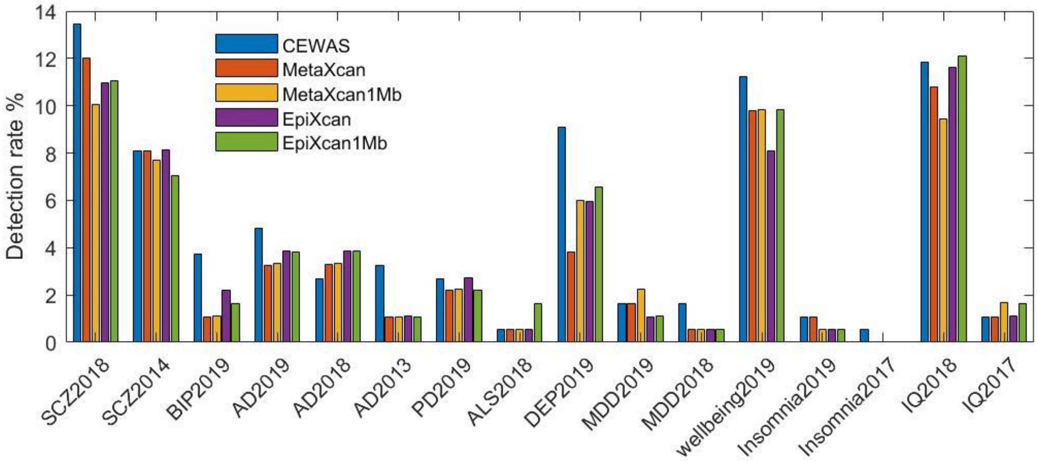


**Fig S1. Detection rate based on distinct signals.** Ratio of the number of significant PCs extracted from the significant genes over the number of significant PCs extracted from the tested genes displayed. The number of significant PCs was used as an estimate of the number of distinct signals within a set of genes (see Methods). This analysis accounts for how some SNPs are shared between models of spatially proximal genes. The overall trend of CEWAS attaining higher detection rate than MetaXcan and EpiXcan remains with this analysis.

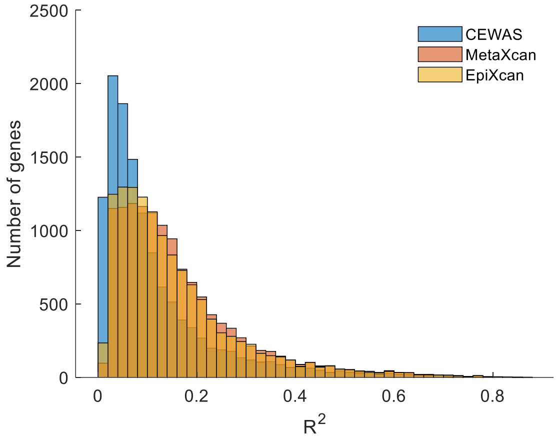


(a) Prediction on ROSMAP data

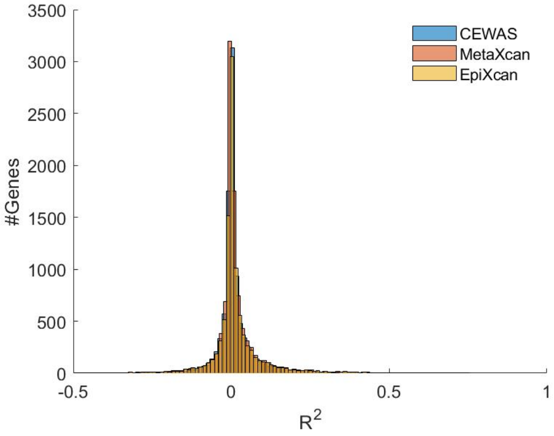


(b) Prediction on CMC data

**Fig S2. $R^2$ of gene models.** (a) $R^2$ estimated by correlating the predicted and measured expression levels in ROSMAP. CEWAS attained lower $R^2$ as expected, since CEWAS is designed to extract a specific component of gene expression, namely the epigenomic component that is driven by genetic effects. In contrast, MetaXcan and EpiXcan are optimized for predicting gene expression. (b) $R^2$ estimated by applying ROSMAP models to CMC genotype data and correlating the predicted expression levels with that measured in CMC. $R^2$ appears similar across methods, but CEWAS actually attained slightly lower $R^2$ if we zoom into the results (see Fig S3).

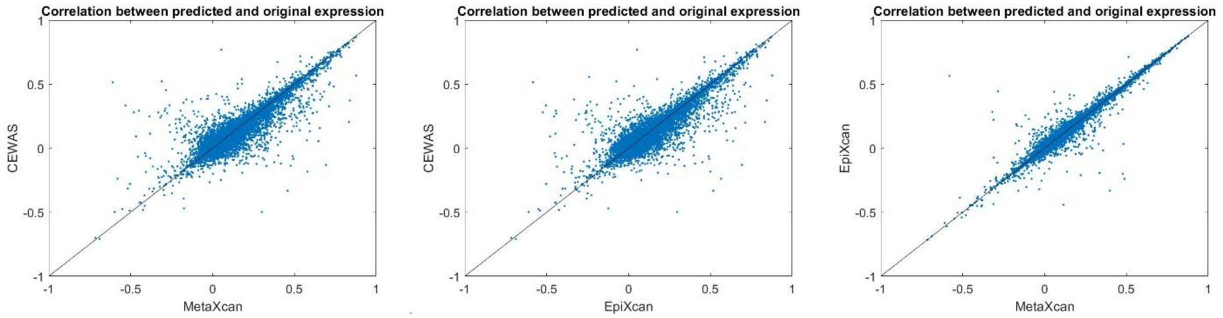


**Fig S3. Correlation between predicted and measured expression levels on CMC data.** Models trained with ROSMAP data were applied to CMC genotype data to predict gene expression. Correlation attained by CEWAS is slightly lower than MetaXcan and EpiXcan.

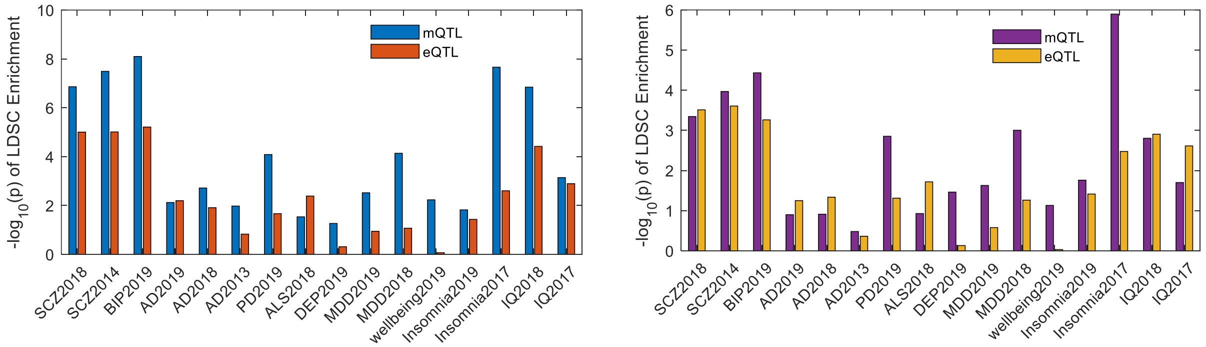


(a) mQTL window = eQTL window = 100Kb

(b) mQTL window = 100Kb, eQTL window = 1Mb

**Fig S4. Stability analysis of partitioned heritability.** Bars show the log p-value of enrichment of mQTLs and eQTLs assessed by LDSC for each GWAS. (a) The enrichment for mQTLs remains higher than for eQTLs with the window size of eQTLs matched to that of mQTLs (p=0.0084, Wilcoxon sign rank test across GWAS). (b) The same trend holds but to a lesser extent when we matched the number of mQTL SNPs to the number of eQTL SNPs by taking mQTL SNPs that are closer to CpGs (p=0.1122, Wilcoxon sign rank test across GWAS; p=0.0269, paired t-test).

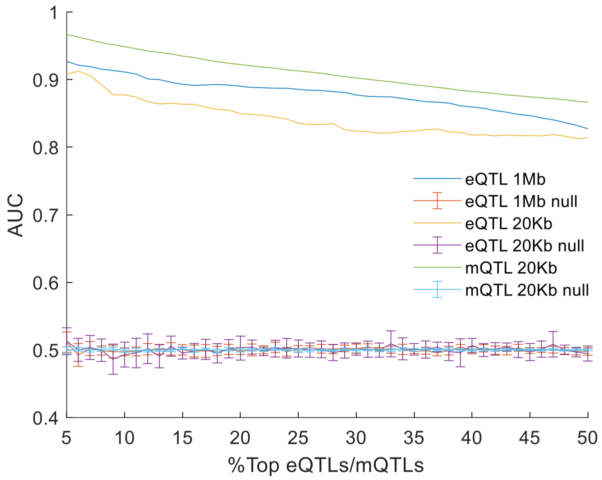


**Fig S5. Replication of mQTLs vs. eQTLs.** Using mQTLs in Jaffe et al.[35] as the reference, we first found the top mQTL SNP for each CpG, and ordered the resulting top mQTLs based on p-values. Taking *q*% of the top mQTLs as "ground truth", *q* = 5% to 50%, we computed AUC on mQTL p-values derived from the ROSMAP data. Since mQTLs from Jaffe et al. were estimated with a 20Kb window, we restricted the ROSMAP mQTLs to those within the same window. The same procedure was applied to estimate AUC for eQTLs, with eQTLs from CMC as the reference. Since eQTLs from CMC were estimated with a 1Mb window, we first examined ROSMAP eQTLs within a 1Mb window, but also examined CMC and ROSMAP eQTLs restricted to the same window size as the mQTLs, i.e. 20Kb. To estimate the null, we permuted the top eQTLs/mQTLs and repeated the procedure.

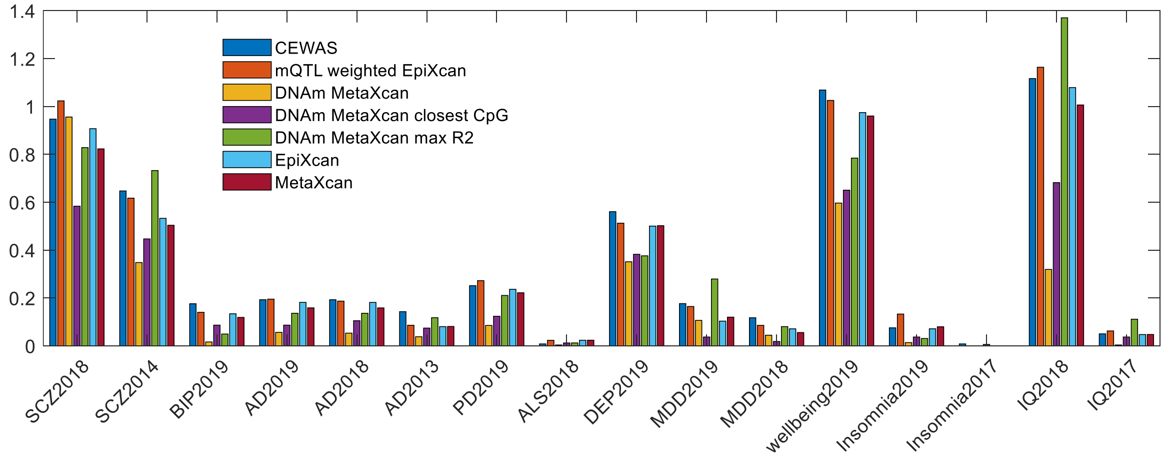


**Fig S6. Gene detection rate with DNAm mediated effects on GWAS phenotypes.** We assessed DNAm mediated effects on GWAS phenotypes in four ways. First, we examined results from models used in the first stage of CEWAS, i.e. MetaXcan models built with each CpG taken as the response and SNPs within ±50Kb from that CpG as predictors. Detection rate of these DNAm MetaXcan models, defined as the number of significant CpGs among tested CpGs, is significantly lower than CEWAS (p=0.0004), MetaXcan (p=0.0013), and EpiXcan (p=0.0006) based on Wilcoxon sign rank test across GWAS. Second, we assessed CpG-to-gene mapping that does not use expression data, by taking the DNAm MetaXcan p-value of the closest CpG of each gene as the p-value of that gene. The gene level detection rate is significantly lower than CEWAS (p=0.0004), MetaXcan (p=0.0003), and EpiXcan (p=0.0004). Third, we took the DNAm MetaXcan p-value of the CpG with the largest $R^2$ in terms of DNAm prediction among CpGs within ±500Kb from each gene as the p-value of that gene. The detection rate is lower than CEWAS (p=0.1331) and EpiXcan (p=0.3808), and higher than MetaXcan (p=0.4235). Fourth, to contrast using epigenomic annotation against explicitly modeling associations between SNPs and CpGs, we modified EpiXcan by weighting the sparse penalty using mQTL p-values. We first associated each CpG to SNPs that are within ±50Kb. We then found the smallest p-value, $p_k$, across CpGs for each SNP $k$, and used $10^{10}p_k+0.5$ (capped at 1) as the weight for its sparse penalty. Multiplying by $10^{10}$ accounts for the number of SNP-CpG pairs tested, and +0.5 penalizes SNPs with the strongest mQTL effects by half the amount as SNPs with weak/no mQTL effects. The detection rate of mQTL-weighted EpiXcan is significantly higher than both MetaXcan (p=0.00006) and EpiXcan (p=0.00006), and lower than CEWAS on average (p=0.3881).

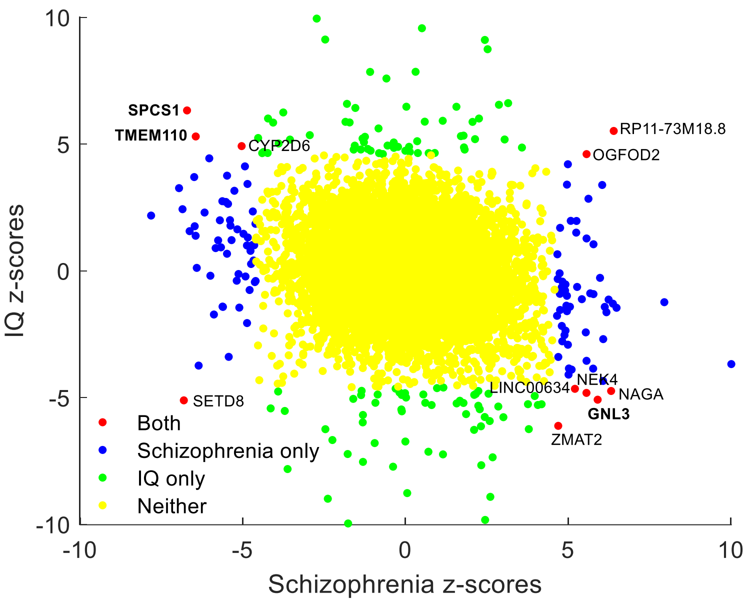


**Fig S7. Genomic correlation between schizophrenia and IQ**. Gene level z-scores estimated by CEWAS for schizophrenia vs. IQ displayed. A genomic correlation of -0.15 was observed. The genes that were detected by CEWAS across schizophrenia, bipolar disorder, depression, and IQ are highlighted in bold. Observing opposite signs in z-scores for the highlighted genes matches how risk alleles in these loci were shown to correlate with lower cognitive test scores[37].

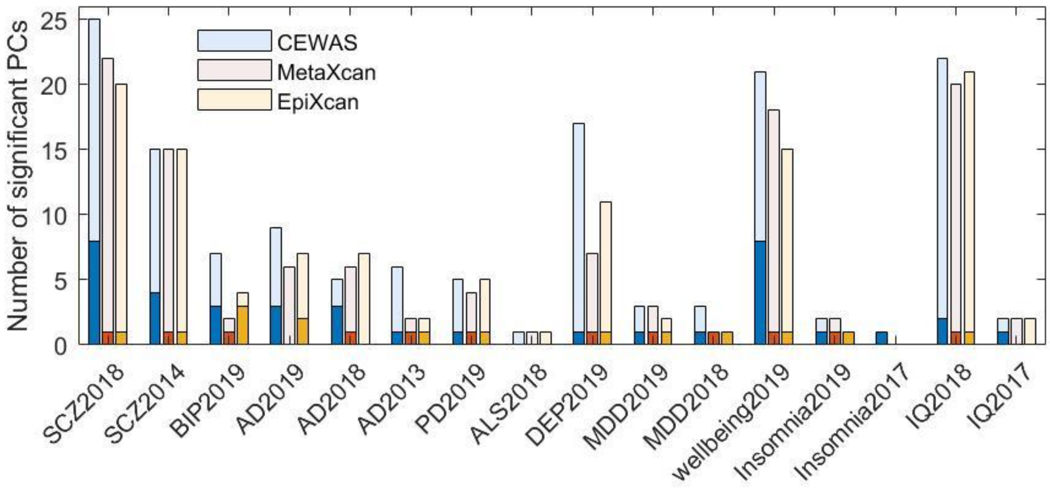


**Fig S8. Number of distinct signals among differential genes.** Number of distinct signals (i.e. number of significant PCs, see Methods) among differential genes (darker shade) and significant genes (lighter shade) detected by each method. Only genes tested in all three methods were considered in extracting differential genes exclusively found by one method but not the other two. The overall trend of CEWAS finding more distinct signals than MetaXcan and EpiXcan remains.

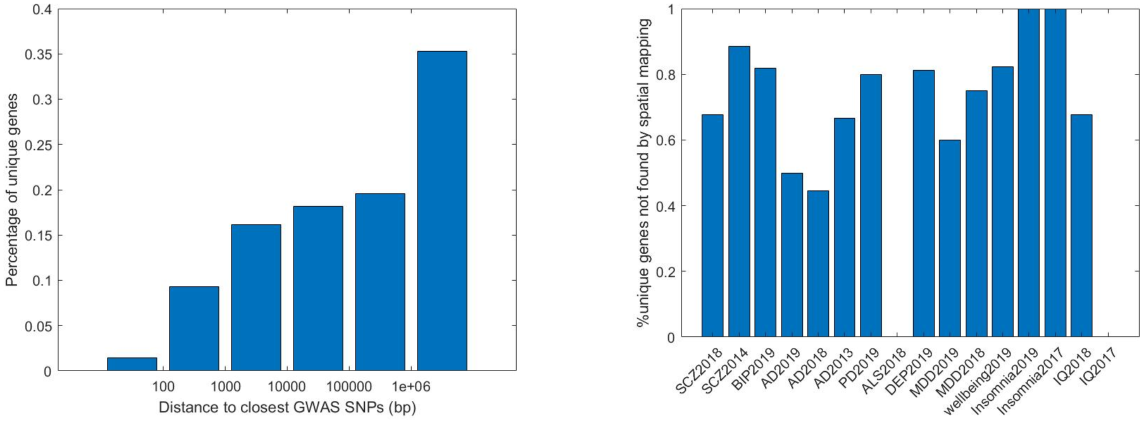


(a) (b)

**Fig S9. Comparison with spatially mapping GWAS SNPs.** (a) The distance between TSS of differential genes exclusively found by CEWAS and their closest GWAS hits summarized. Each bar corresponds to the percentage of differential genes lying in a range of distance, e.g. a bar between 100 and 1000 corresponds to the percentage of differential genes with distance between 100 and 1000 base pairs. More than half of the differential genes are >100Kb away from any GWAS hits. (b) The percentage of differential genes exclusively found by CEWAS but missed by spatially mapping GWAS SNPs to their closest genes shown. Note that no differential genes were found for ALS2018, hence why the percentage is zero.

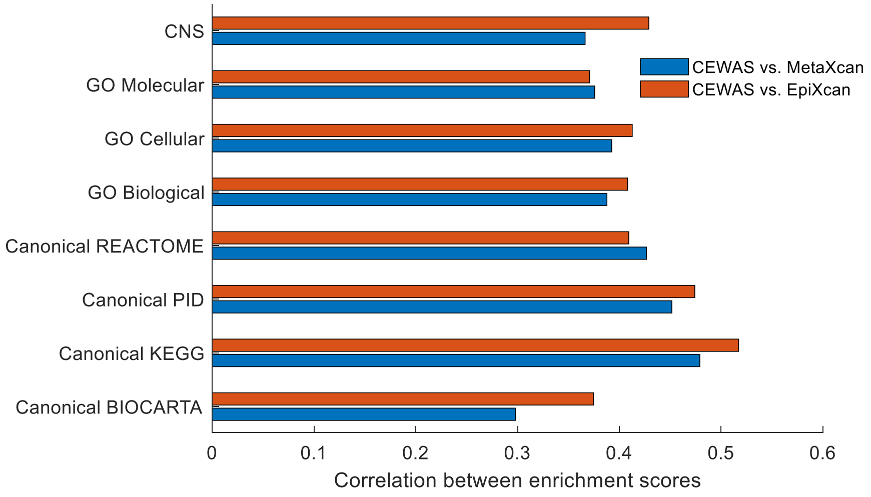


**Fig S10. Correlation of gene set enrichment scores between CEWAS and contrasted methods.** The average correlation over GWAS for each gene set category displayed. The observed correlation suggests moderate similarity in enriched gene sets between CEWAS and the contrasted methods, which we confirmed by manual inspection.

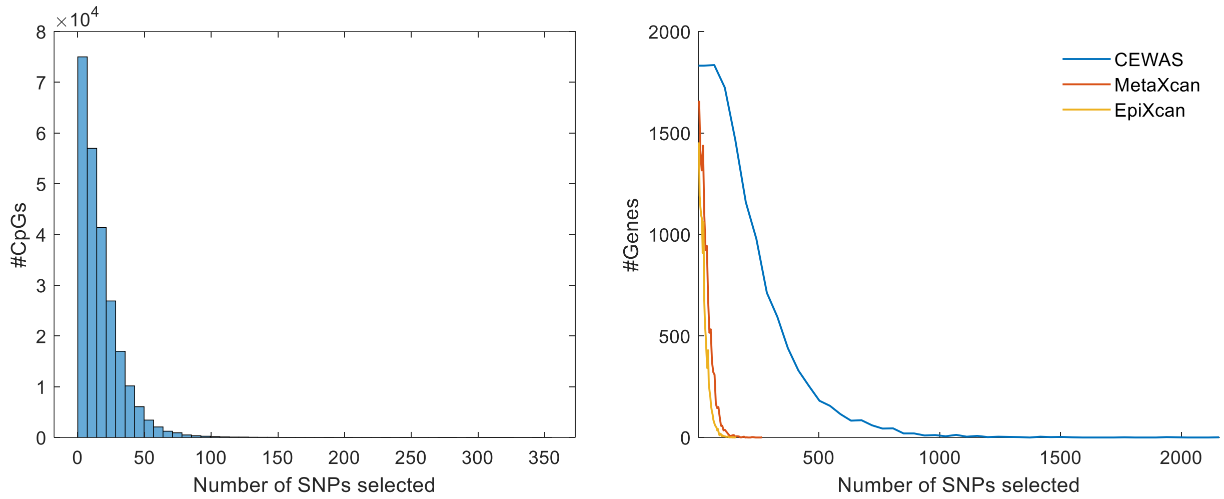


(a) Number of SNPs selected per CpG

(b) Number of SNPs selected per gene.

**Fig S11. Number of selected SNPs.** (a) Number of SNPs selected per CpG by elastic net. (b) Number of SNP selected per gene by elastic net. CEWAS tends to have more SNPs selected since 543 samples are available per CpG for selecting relevant SNPs with elastic net and each gene is associated with 9.62 CpGs on average, whereas MetaXcan and EpiXcan have only 534 samples per gene for SNP selection.
